# The effect of outdoor recreation on mammal habitat use and diversity revealed by COVID-19 closures

**DOI:** 10.64898/2026.04.02.715670

**Authors:** Alexandra Dimitriou, Sarah Benson-Amram, Kaitlyn M. Gaynor, A. Cole Burton

## Abstract

The rising demand for outdoor recreation worldwide may be undermining the conservation objectives of protected areas (PAs). We leveraged a natural experiment, in which two adjacent PAs were closed to the public for different durations during the COVID-19 pandemic. Using detections from 39 camera traps in Joffre Lakes and Garibaldi Parks, Canada, from 2020-2022, we examined how recreation influenced mammal habitat use and diversity. Bayesian regression showed weak evidence that, when recreation was higher, detections declined for black bear, mule deer, and marten, while detections of bobcat and hoary marmot shifted closer to trails. Accumulation curves revealed that species richness and diversity were higher in the closed vs. open PA in 2020 (mean differences of −5.04 for richness and −0.33 for Shannon diversity). However, diversity did not decline consistently despite increases in recreation in 2021 and 2022. Notably, several rare species were only detected in the lower-recreation PA, suggesting they may be filtered out of the higher-recreation PA. This emphasizes the need for long-term monitoring to detect delayed and cumulative effects of recreation on mammal communities. Given growing global pressures on biodiversity, we urge PA managers to prioritize adaptive management to assess and balance outdoor recreation with conservation goals.

## 1. Introduction

The Anthropocene has ushered in an era of global biodiversity loss (Butchart et al., 2010; Johnson et al., 2017). Amid rapid land use change, protected areas (PAs) are a prominent conservation strategy to preserve ecosystems from anthropogenic pressures and foster ecosystem resilience (Berkes et al., 2008; Chen et al., 2022; Pacifici et al., 2020; Watson et al., 2014). PAs also play an important role for human communities by boosting local economies through nature-based tourism and supporting the mental and physical health of visitors through outdoor recreation (Bushell & McCool, 2007; Winter et al., 2020). In recent years, demand for outdoor recreation has grown considerably, and accordingly, so have concerns about the increasing pressure on protected ecosystems (Beery et al., 2021; Schulze et al., 2018; Weststrate et al., 2025). Despite the dual mandate of many PAs to balance recreation and conservation, it is becoming evident that most PA managers lack the data needed to understand whether current or future levels of recreation will compromise conservation goals (Geldmann et al., 2013; Jones et al., 2018; Reed & Merenlender, 2008).

The growing field of recreation ecology aims to uncover the complex impacts of outdoor recreation on natural systems, including wildlife. Researchers have reported a range of behavioural and physiological responses to recreation by wildlife, such as spatiotemporal avoidance of recreationists (Anderson et al., 2023; Gaynor, Hayes, et al., 2025; Naidoo & Burton, 2020; Salvatori et al., 2023; Thompson et al., 2025), changes to vigilance behaviours (Mainini et al., 1993; Naylor et al., 2009), physiological stress (Arlettaz et al., 2007; Creel et al., 2002; Piñeiro et al., 2012), and reductions in fitness, body condition, reproductive success, and parental investment (Finney et al., 2005; Frid & Dill, 2002; Wilson et al., 2020). On the other hand, many studies show minimal or variable associations between wildlife and recreation, with few studies investigating whether individual- and population-level responses translate into community-level changes in species diversity and composition (Bro-Jørgensen et al., 2019; Larson et al., 2016; Smith et al., 2024). This is an important gap to fill, as the restructuring of communities in response to anthropogenic disturbances can have important impacts on ecological functions and services (Kunze et al., 2025; Ordiz et al., 2021; Wilson et al., 2020).

Conserving a diversity of species and functional roles is critical for maintaining ecosystem stability; however, wildlife species are differentially affected by anthropogenic disturbances (Duffy, 2009; Lacher et al., 2019; Sévêque et al., 2020). In mammals, larger-bodied, longer-living, and more carnivorous species tend to be more vulnerable, as their dietary needs and large home ranges increase their likelihood of conflict with humans, sometimes leading to their exclusion from human-dominated landscapes (Baker & Leberg, 2018; Hill et al., 2020; Reed & Merenlender, 2008; Suraci et al., 2021). Our understanding of human-wildlife interactions is also informed by the landscape of fear framework, which describes an animal’s perception of its environment based on a cost-benefit analysis associated with predation risk (Gaynor et al., 2019; Laundré et al., 2001). This framework has been applied to describe the way that human presence (even when engaging in non-consumptive recreational activities) may exert spatially and temporally explicit patterns of perceived risk on wildlife, motivating them to alter their behaviour to avoid interacting with humans (Frid & Dill, 2002; Gaynor et al., 2019; Palmer et al., 2023). Large predators may be more likely to be displaced by fear of humans, due to the severity of potential conflict, and this displacement may in turn create refuge for prey species and subordinate predators, a phenomenon termed the human shield hypothesis (Berger, 2007; Gaynor, Wooster, et al., 2025; Granados et al., 2023; Prugh et al., 2023; Ritchie & Johnson, 2009). By displacing large predators and altering risk landscapes, human presence acts as a behavioural filter with cascading effects on species interactions and community structure.

Species traits can thus shape their capacity to navigate and persist in human-altered environments, and differential responses across species can in turn affect community-level outcomes. Species with faster life histories (rapid maturation, high fecundity, short lifespans, reduced parental investment) may be more capable of persisting in risky environments; they also tend to have smaller body sizes and home ranges, which lowers their likelihood of conflict with humans (Hill et al., 2020; McKinney & Lockwood, 1999; Suraci et al., 2021). Species with greater niche breadth have also been reported to succeed in human-dominated landscapes because of their flexibility in habitat selection and their success in exploiting novel anthropogenic food sources (Barrett et al., 2019; Newsome & Van Eeden, 2017; Nickel et al., 2020; Santini et al., 2019). An understanding of the role of species traits in mediating the effects of recreation on wildlife can shed light on how human disturbance in protected areas might filter and disrupt ecological communities (Smith et al., 2024).

A key challenge in understanding the impacts of recreation on wildlife is disentangling its effects from the many other factors that shape wildlife behaviour and often covary with recreation, such as seasonal variation in environmental resources and conditions. During the COVID-19 pandemic “Anthropause,” the significant reduction (or even absence) of recreation in PAs provided a rare opportunity to isolate the effects of recreation on wildlife activity by creating a quasi-experimental control (Anderson et al., 2023; Bates et al., 2020; Burton et al., 2024; Rutz et al., 2020). Comparing wildlife activity during this “low recreation control” with periods of typical or even elevated recreation offers valuable insights that could inform PA management (Gaynor, Hayes, et al., 2025; Rutz, 2022).

In this study, we used camera trap detections to examine responses of 15 terrestrial mammal species (Table S1) to changes in recreation during and after COVID-19 closures of two highly visited PAs. We hypothesized that higher recreation would negatively affect large mammal species, with species functional traits shaping interspecific variation in sensitivity to recreation (i.e., the degree to which they avoid recreationists; Table 1). We expected these differential responses to result in a negative association between recreation and species richness and community evenness. Under conditions of high recreation, tolerant species would be more dominant and sensitive species would be displaced, leading to a simpler and more homogenous mammal community (Grimm et al., 2008; McKinney & Lockwood, 1999; Newbold et al., 2018). To test these hypotheses, we examined how monthly recreation intensity and proximity to hiking trails influenced large mammal detections from camera traps across multiple years, including the 2020 closure of one PA, and compared diversity metrics across years.

**Table 1.** Hypothesized traits associated with mammal responses to recreation, as derived from the literature (further citations provided in main text). Species-specific trait values are listed in Table S1, and trait-based sensitivity scores in Table S2.

| Species trait | Predicted relationship with recreation | References |
| --- | --- | --- |
| Body mass | Larger-bodied species are more sensitive | (Salvatori et al., 2023; Vargas Soto et al., 2022) |
| Trophic level | More carnivorous species are more sensitive | (Baker & Leberg, 2018; Ripple et al., 2014; Suraci et al., 2019) |
| Niche breadth | Species with more limited habitat and diet breadth are more sensitive | (Nickel et al., 2020; Santini et al., 2019) |
| Generation length | Species with longer generations (proxy for slower life history strategy) are more sensitive | (Hill et al., 2020; McKinney & Lockwood, 1999) |
| Dispersal | Wider ranging species are more sensitive | (Crooks, 2002) |

## 2. Methods

### 2.1 Study Area

We conducted this study on and around hiking trails in two of the most visited PAs in southwestern British Columbia, Canada: Garibaldi Park and Pipi7íyekw/Joffre Lakes Park (henceforth referred to as “Garibaldi” and “Joffre Lakes”; Figure 1). These two parks, located in the Coast Mountains, are very popular destinations for hikers in and around the nearby Vancouver metropolitan area, receiving 190,000 and 160,000 annual visitors, respectively (BC Parks, n.d.). In the last decade, both parks have seen a considerable surge in visitation, which can largely be attributed to their popularity on social media (Baluja, 2016; Bergman et al., 2022; Kohlhardt et al., 2018). Between 2010 and 2019, the smaller (15 km^2^) Joffre Lakes saw a 222% increase in park visitation (Ministry of Environment and Climate Change Strategy, 2021).

**Figure 1.**
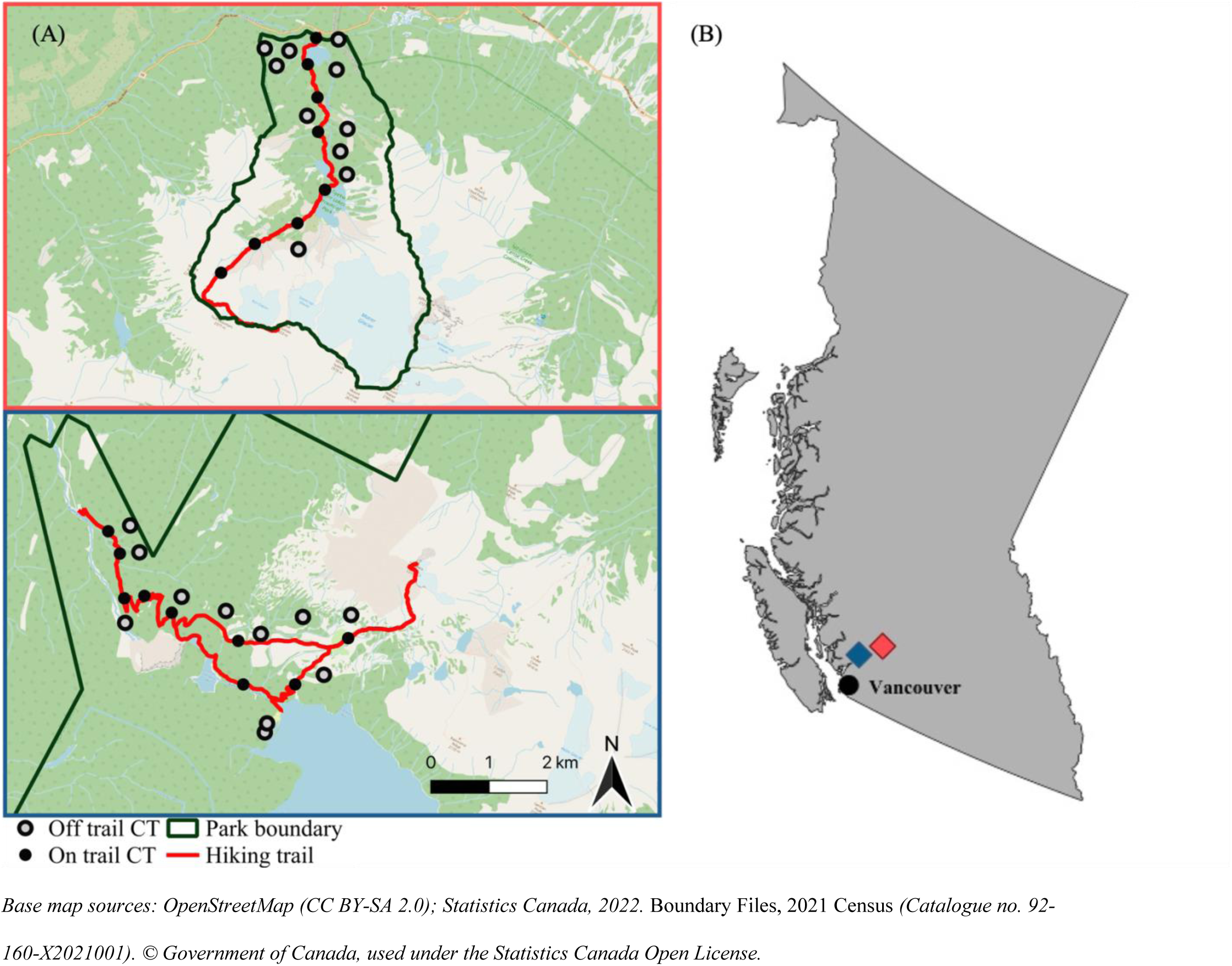
Study area of Joffre Lakes Park (red) and Garibaldi Park (blue) (A) in southwestern British Columbia, Canada (B). The provincial park boundaries are represented by the dark green outlines. Hiking trails of interest are represented by red lines. Camera traps (CT) are represented by black points (on trail) and grey points (off trail). In panel A, green represents forested areas, yellow lines represent highways, blue represents bodies of water, light blue represents glaciers, white and beige represent bare rock and snow in high alpine areas.

Garibaldi is a much larger park (1950 km^2^); however, we focused on a small section of the park (Rubble Creek to Garibaldi Lake) which receives the most recreation and has comparable features to the trail of interest in Joffre Lakes (e.g. a gradual ascent through mountainous terrain to a scenic glacier-fed lake). Dogs and motorized vehicles are not permitted in either PA, and bicycles are not permitted in Joffre Lakes or in the Garibaldi study area; thus, the main recreation activity is hiking. The two PAs offer very similar seasonal, environmental and topographic conditions, and the expected mammal community compositions are very similar based on published ranges; therefore, we treated the two areas as replicate sampling areas for comparison (Figure S1; Table S3; IUCN, 2024).

In response to the COVID-19 pandemic, both PAs were closed to the public but for different durations. Joffre Lakes was closed for 14.5 months (April 8^th^, 2020 - June 22^nd^, 2021), while Garibaldi was only closed for about 3.5 months (April 8^th^, 2020 - July 27^th^, 2020), reopening just before our study began in August 2020. Although data were collected year-round, we restricted our comparisons to the snow-free months of each year (May to October), as species occurrence varies seasonally due to factors such as hibernation/torpor (e.g. bears) and seasonal elevational migrations (e.g. mule deer; Geiser, 2013; Monteith et al., 2011). Hiking in these parks is also concentrated during the snow-free months (Figure S3).

Based on these quasi-experimental differences in recreation across parks and time periods, we developed a set of predictions. We considered that the first period of sampling (2020) in Joffre Lakes would represent a “no recreation” control and thus predicted it to have the greatest species richness and community evenness, as more sensitive species would use the trail in the absence of recreation. We expected the 2020 patterns in Joffre Lakes to contrast with those in Garibaldi, which was open. We predicted Garibaldi to have lower richness and evenness than Joffre Lakes in 2020 due to less use by sensitive species and more use by more tolerant species. In subsequent years (2021 and 2022), both PAs were open and thus we expected similar levels of recreation. We therefore predicted similar patterns of diversity metrics and habitat use (i.e., richness, evenness, and detections of sensitive species would decrease from 2020 to 2021 in Joffre Lakes but remain stable in Garibaldi).

### 2.2 Camera Trap Survey

Camera traps (CTs) are a reliable, cost-effective, and non-invasive tool used to sample multiple species simultaneously and over long periods without requiring direct observation (Burton et al., 2015; Kays et al., 2009; Wearn & Glover-Kapfer, 2019). CT sampling has also been extended to quantify recreational activity (Fennell et al., 2023; Naidoo & Burton, 2020; Procko et al., 2022). In August 2020, we deployed 18 CTs in Joffre Lakes and 21 CTs in Garibaldi (Figure 1). The cameras were active for two years from August 2020 to September 2022. We used Reconyx HyperFire2 (Reconyx, Holmen, USA) CTs fastened to trees, approximately 0.5–1 m off the ground, facing perpendicular to a target area of expected animal or human travel (i.e., hiking or game trail) at a distance of 3-5 m. CT locations were chosen randomly within two strata in each park: on and off of hiking trails. CTs were deployed at least 250 meters apart, with off-trail CTs positioned between 145 and 687 meters from the nearest hiking trail. The CTs were programmed to collect motion-triggered images as well as a single time-lapse image each day at noon to monitor functionality.

### 2.3 Data Processing

After collection, CT images were processed and identified to species level using a custom database. We used Megadetector, an open-source object detection model, to separate images of humans, animals, and vehicles for identification (Beery et al., 2019; Fennell et al., 2022; Mitterwallner et al., 2024). Photos that contained human images were subsequently blurred to protect the privacy of park users. All CT images identified as focal species were subjected to a secondary quality check by a senior member of the research team (Silva-Rodríguez et al., 2025). CT images of especially rare species are included in the Supporting Information (Figure S2).

Prior to modelling, we set a threshold of independence of 30 minutes for detections of wildlife species and 1 minute for detections of humans at each CT, as is common practice in similar studies (Burton et al., 2015, 2024; Procko et al., 2022). We assumed that detections of a single species at a given CT site were the same individual or part of a non-independent group (e.g. a family unit) if consecutive detections were within 30 minutes. During typical periods of recreation activity, humans were more continuously detected at CTs, such that many different individuals and groups would be detected during the 30-minute threshold. We, therefore, used the shorter 1-minute interval and assumed that detections of humans captured after a period without detections of at least 1 minute were likely to be different individuals or groups.

For statistical analysis, we only used detections of mammal species with an average adult body mass greater than 250g (and thus were likely to be more reliably detected by CTs; Table S1). For the habitat use models, we selected focal species as those which were detected in both PAs, with a minimum of 35 total independent detections over the entire study period (i.e., exposed to the full range of recreation intensity, including the control period, and sufficient sample size for modelling; Table S1; Kays et al., 2020)). For the community models, all species that met the body mass threshold were included, regardless of the number of detections.

### 2.4 Modelling habitat use

We used Bayesian generalized linear mixed effects models (GLMMs) of CT detection rates to assess habitat use by focal species across spatial and temporal variation in recreation intensity. Rather than treat finer-scale temporal variation in detections as a source of error, such as in an occupancy modelling framework (Neilson et al., 2018), we treated this variation as a signal of the ecological process of interest (i.e., animal abundance and activity at a site). Our response variable was the number of independent detections per month for each species, which we interpreted as an index of habitat use by one or more individuals of the same species in the area around the CT site (Fennell et al., 2023; Tattersall et al., 2020). We used a negative binomial distribution to account for overdispersion of the detection counts. In addition to models for each of the focal species with at least 35 independent detections, we ran a rare carnivore model which aggregated five carnivore species (Canada lynx, cougar, grey wolf, grizzly bear and wolverine) with insufficient detections to support single-species models (Table S2). All models included a random effect for CT site to account for non-independence among repeated monthly counts of detections at individual CT sites. We tested for spatial autocorrelation of residuals using Moran’s I tests, which showed limited spatial dependence across species and months (Figure S4). We also tested for temporal autocorrelation of residuals using Durbin-Watson tests, which indicated that autocorrelation was within acceptable limits (i.e., 1.50-2.50; Table S4; Durbin & Watson, 1950)

In calculating our recreation and habitat use variables, we defined a month as a 28-day period between May and October, starting from Tuesday until Wednesday, to ensure that each “month” contained exactly 4 full weekends, placed centrally within the month, as recreation intensity is greatest during weekends (Fennell et al., 2023; Green et al., 2023; Longshore et al., 2013). We included an effort offset term (log CT days) in our models to account for any periods of camera inactivity within a month and excluded the final month of the study (August 24^th^ – September 20^th^, 2022), in which cameras were only active 1-7 days. We chose this monthly time scale to avoid an excess of repeats with zero species detections while still assessing responses to the temporally variable recreation intensity. Our primary predictor variables of interest were the effort-corrected Park-Month recreation intensity (number of independent human detections across all CTs per month per 100 CT days; hereafter, recreation intensity), as a measure of temporal recreation activity, and an interaction between recreation and proximity to trail, as a measure of spatial recreation activity. We initially considered three scales of recreation intensity which could influence habitat use, ranging from a coarse Park-Year scale (human detections from May-October of each year), Park-Month, and the localized Park-Site (Figure 2). We chose the intermediate Park-Month metric to capture sufficient variation in recreation intensity while accounting for the broader zone of influence that on-trail recreation may exert on off-trail sites, under the assumption that human activity can impact habitat use beyond the immediate vicinity of trails (Thompson et al., 2025). Given that both focal trails are less than 10 km in length, and most hikers complete the full trail, we assumed relatively uniform recreation pressure across on-trail sites in a given month. In addition to serving as concentrated areas of recreation activity, trails may also function as travel corridors for wildlife, facilitating unobstructed movement along linear features (Dickie et al., 2020; Fitzpatrick et al., 2024). Thus, we included a proximity to trail variable with an exponential decay function (*e*^−*distance to trail*/500*m*^), which assumes that the influence of trails rapidly declines with distance, particularly within 500 meters. To assess whether the effect of monthly recreation was moderated by proximity to trail, we included an interaction term between recreation intensity and proximity to trail. Including both spatial and temporal recreation variables, along with their interaction, allowed us to assess species’ responses to both direct and indirect forms of recreational pressure.

**Figure 2.**
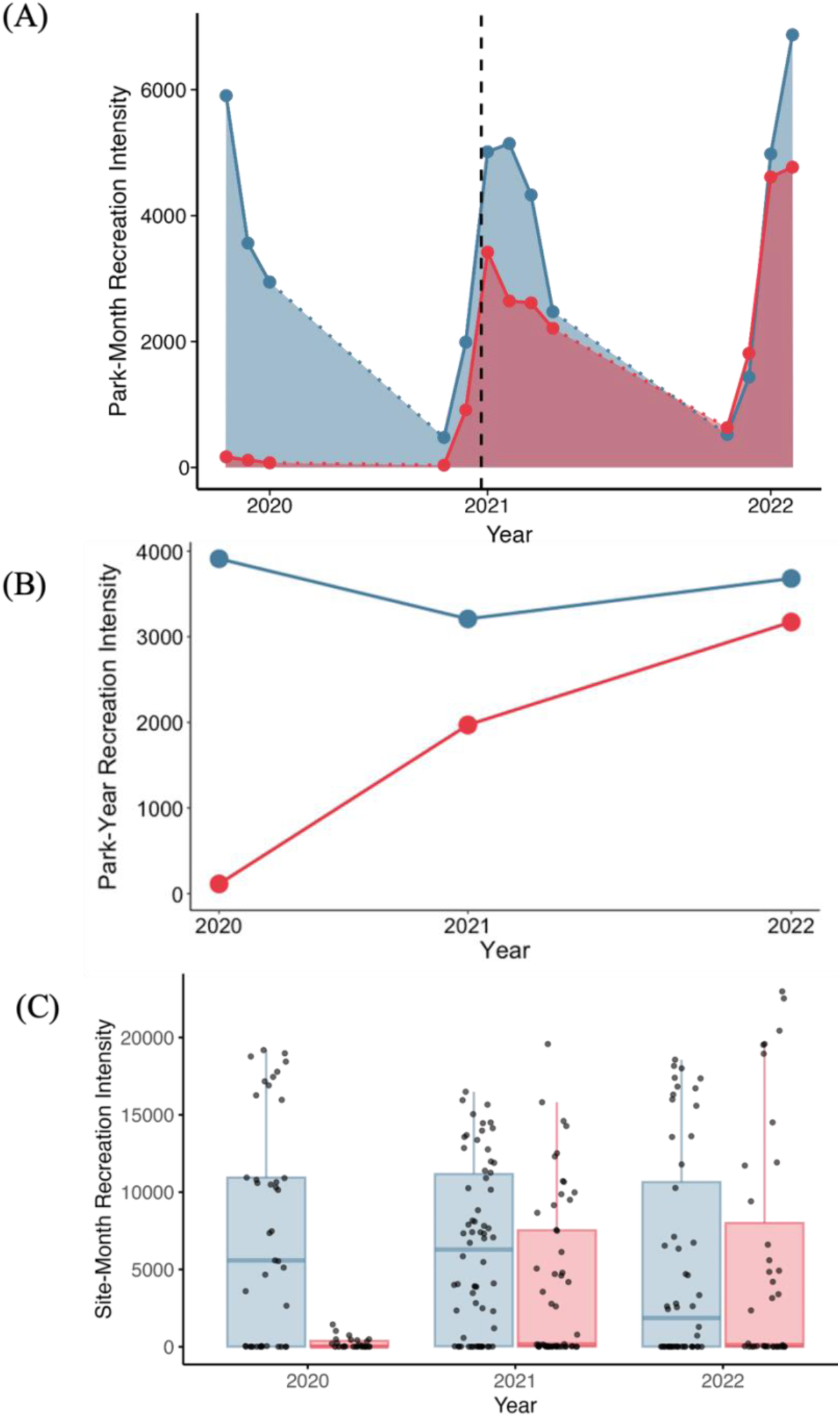
Recreation intensity in Joffre Lakes (red) and Garibaldi (blue) during the study period at three spatiotemporal scales. A) Park-Month recreation intensity: number of independent human detections summed across all CTs in each park per month, standardized per 100 CT days. Dotted segments indicate periods outside the study months that were excluded from habitat use models (Figure S3 includes all months that the CTs were active). The black dashed vertical line marks the reopening of Joffre Lakes on June 22, 2021. (B) Park-Year recreation intensity: number of independent human detections per park and year (for months included in the habitat use models), standardized per 100 CT days. (C) Site-Month recreation intensity: number of independent human detections per CT site, per month, standardized per 100 CT days, shown by park and year. Boxes span the interquartile range; horizontal lines represent the median. Datapoints represent raw values; those beyond the whiskers are considered outliers.

We also included additional predictor variables to control for spatial and temporal variation in environmental conditions that we anticipated would influence species habitat use (Table 2). Specifically, we included an index of primary productivity (Normalized Difference Vegetation Index, NDVI, from MODIS13Q1, summarized by overlap with 28-day survey months), which we assumed to reflect variation in forage availability and seasonality (Pettorelli et al., 2005, 2011; Tuck et al., 2014). We included canopy cover to account for habitat structure, measured as the percentage of vegetation above 2 m from the National Terrestrial Ecosystem Monitoring System (NTEMS; Matasci et al., 2018). Canopy cover was strongly correlated with elevation (r=0.59; Figure S5). This system is a mosaic of contrasting high alpine tundra and dense coastal cedar-hemlock rainforest characterized by tall trees and patchy canopy openings that support understory growth (Figure 1; Weetman & Prescott, 2001). We therefore assume that overstory and understudy density are linked, providing concealment cover to smaller and larger mammals alike (Camp et al., 2013; Comer et al., 2025; Potash et al., 2019). All predictor variables were assessed for collinearity using Pearson’s correlation coefficient and only variables with pairwise correlations less than |0.5| were included in the final models (Figure S5). Subsequently, numerical predictor variables were standardized by subtracting the mean and dividing by the standard deviation (except the decay-transformed proximity to trail, which was already bounded between 0-1).

**Table 2.**
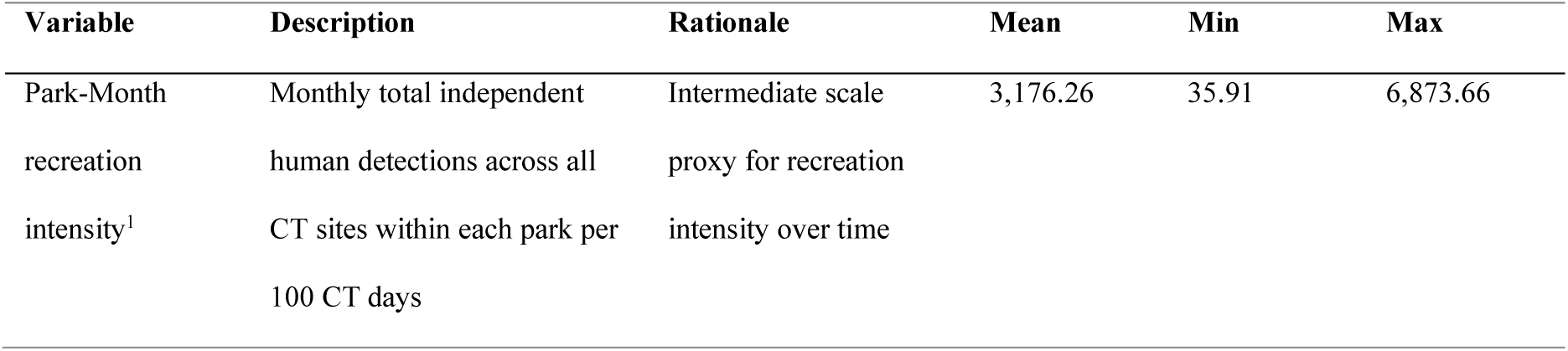
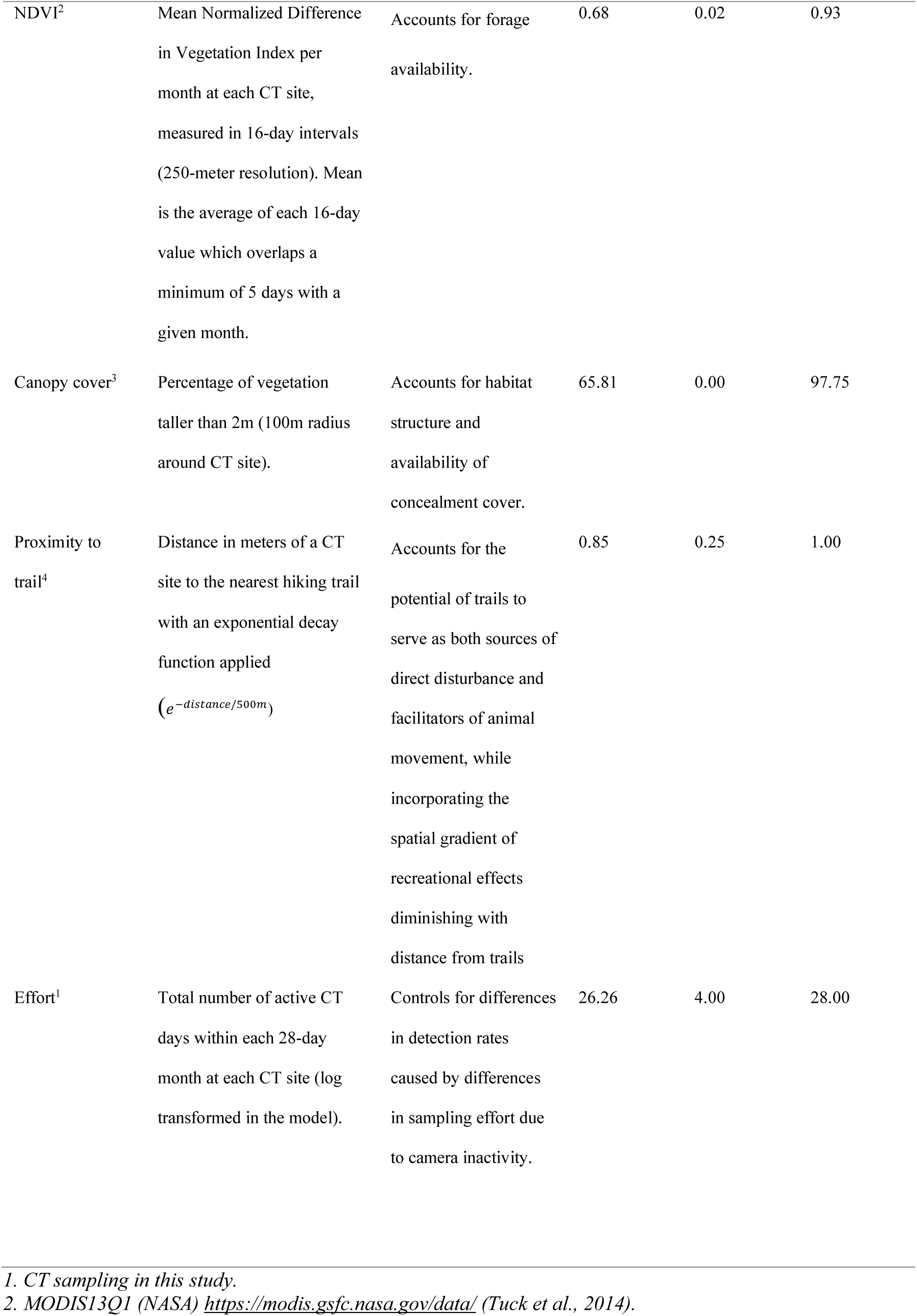
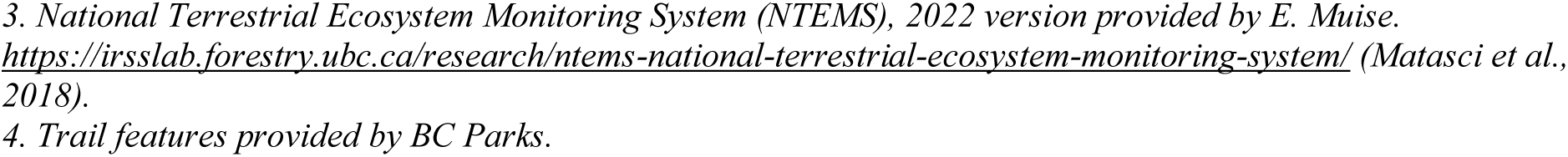
Predictor variables used in the Bayesian GLMMs to investigate the effect of recreation intensity on monthly habitat use for focal species, measured by camera trap (CT) sampling in Joffre Lakes and Garibaldi Parks. Summary values (mean, min, max) were calculated across all 39 CT sites deployed across both parks, and throughout the months included in habitat use models July 29^th^- October 20^th^, 2020; May 5^th^- October 19^th^, 2021; May 4^th^ – August 23^rd^, 2022).

We implemented the GLMMs using the *brms* package in R (Bürkner, 2017; R Core Team, 2024), using weakly informative priors. We specified a normal prior, centred around 0, with a standard deviation of 2 for the fixed effects and intercept, which provides mild regularization while still allowing sufficient variation to detect meaningful effects (Gelman et al., 2013). We specified a Cauchy prior centred around 0 with a scale parameter of 2 on the standard deviation of the random effect to allow for large variability between CT sites while avoiding extreme values (Gelman, 2006). For all species, we ran the models with 4 chains of 10,000 iterations with a burn-in period of 4,000, a thin rate of 1, an adapt delta of 0.999 and a maximum tree depth of 20 to improve model stability. We ran each species model with the full set of predictor variables (Table 2). We assessed model convergence based on Rubin-Gelman statistics (Rhat < 1.10), effective sample size (>1000), inspection of trace plots and posterior predictive checks. We considered parameter estimates with 95% Bayesian credible intervals drawn from the posterior distribution that do not overlap zero as strong evidence of an association between a predictor variable and species’ habitat use, and 80% credible intervals that do not overlap zero as weak evidence of an association (Fennell et al., 2023; Granados et al., 2023; Kéry & Royle, 2020).

### 2.5 Modelling Community Shifts

We assessed community-level responses to recreation by comparing species richness, Shannon diversity and Pielou’s evenness between the two PAs in three distinct periods: summer-fall of 2020, 2021, and 2022. Species richness, the number of unique species present at a given location, is a fundamental and simple measure of biodiversity often used to assess the effects of disturbance on wildlife communities and evaluate management actions (Chen et al., 2022; DeWan & Zipkin, 2010; Gwinn et al., 2016). However, species richness alone does not capture more subtle changes in community structure when the species pool remains unchanged but relative abundances of species change (Hillebrand et al., 2018; Vellend, 2017). Shannon diversity complements richness by incorporating species’ relative abundances (Roswell et al., 2021). Together, these metrics allow us to estimate community evenness, providing a broad understanding of shifts in community structure.

Assessing community composition using CT surveys requires sufficient sampling effort and addressing imperfect detection (Kays et al., 2020; Kéry & Schmidt, 2008; Rovero & Zimmerman, 2016; Rowcliffe et al., 2008). Chao estimators are commonly used to estimate species richness and diversity by accounting for undetected species based on the occurrence of rare species and standardizing estimates by sample completeness (Chao et al., 2014). We derived estimates of richness and diversity from species accumulation curves using the *iNEXT* package in *R* (Chao et al., 2014; Hsieh et al., 2016; S3). We compared species accumulation curves between parks for a 90-day period from the spring/summer to the fall of each year. The 2020 period (August 6^th^, 2020- November 3^rd^,2020) was during the closure in Joffre Lakes and following the re-opening of Garibaldi. The 2021 period (June 22^nd^, 2021- September 19^th^, 2021) began the day that Joffre Lakes reopened to the public, and the 2022 period (May 10^th^, 2022-August 7^th^, 2022) was approximately one year later. For each curve, we extracted the asymptotic estimates of species richness and diversity. Then, we used the estimates of richness and diversity to calculate Pielou’s evenness index as a relative measure of community evenness (Jost, 2010; Pielou, 1966). Specifically, diversity was log-transformed to obtain Shannon entropy and then divided by the logarithm of estimated species richness. Confidence intervals for evenness were calculated using first-order Taylor expansion error propagation, based on the uncertainty of the *iNEXT* estimates.

Finally, we used bootstrap hypothesis tests (with 10,000 replicates) to perform pairwise comparisons of diversity measures between periods within each park, and between parks within each period (Table S8). We used this approach because it enables hypothesis testing and uncertainty estimation directly from model-derived point estimates (e.g., from *iNEXT* curve asymptotes) and their associated standard errors (Efron & Tibshirani, 1994). To avoid zero p-values, we applied a continuity correction ((r+1) / (B+1)); Hall, 1987). To account for multiple comparisons and potential dependencies arising from comparisons across both parks and years, we applied a Benjamini–Hochberg correction (Benjamini & Hochberg, 1995; Table S5).

## 3. Results

### 3.1 Summary of CT survey

The 39 CTs deployed across Joffre Lakes and Garibaldi collected a total of 3,727,244 motion-triggered images over 25,762 CT days (Figure S6). We detected 15 medium-large sized mammal species, with the highest number of independent detections of snowshoe hare (*Lepus americanus*, n = 859), mule deer (*Odocoileus hemionus*, n = 730), and American black bear (*Ursus americanus*, n = 249), and the fewest detections of grizzly bear (*Ursus arctos*, n = 1), Canada lynx (*Lynx canadensis*, n = 1) and raccoon (*Procyon lotor*, n = 4). Seven species met our sample size criteria for the habitat use models: black bear, American marten (*Martes americana*, n = 145), snowshoe hare, bobcat (*Lynx rufus,* n = 42), coyote (*Canis latrans*, n =35), hoary marmot (*Marmota caligata*, n=48), and mule deer (Table S1). Our aggregated rare carnivore model included 30 combined detections of grey wolf (*Canis lupus*, n = 5), grizzly bear, wolverine (*Gulo gulo*, n = 8), cougar (*Puma concolor*, n = 15) and Canada lynx.

Recreation varied considerably between parks and over time. Park-Month recreation intensity varied from a low of 36 independent human detections per 100 CT days in Joffre Lakes during spring 2021 to a high of 6,874 in Garibaldi during summer 2022 (Table 2; Figure 2A). On average, Garibaldi had greater Park-Month recreation intensity than Joffre Lakes (3,855 vs. 2,277). At the Park-Year scale, Garibaldi showed relatively consistent recreation intensity across years (3,911 in 2020, 3,208 in 2021, and 3,681 in 2022), while Joffre Lakes showed a marked increase over time, from only 113 during the closure (2020) to 1,970 in early reopening months (2021) to 3,171 one year after reopening (2022) (Figure 2B). Site-Month recreation intensity also varied considerably, from 4 independent human detections per 100 CT days at an off-trail CT site in Garibaldi to 22,979 independent human detections per 100 CT days at an on-trail CT site in Joffre Lakes in 2022 (Figure 2C).

### 3.2 Drivers of wildlife habitat use

Three of the eight species modelled (including the rare carnivore group) exhibited weak evidence of a negative association between habitat use and monthly recreation intensity, while two exhibited weak evidence for a positive and interactive effect of recreation intensity and trail proximity. Specifically, black bear, mule deer, and marten were detected less in high recreation months, whereas bobcat and marmot were detected closer to hiking trails under high recreation (Figure 3). Regardless of recreation intensity, we also observed weak evidence of a positive association between detection rates and proximity to trail for martens. Among environmental variables, canopy cover had the greatest association with habitat use, being strongly negative for marten and marmot, and strongly positive for hare. There was strong positive association with NDVI for deer, weak positive association for black bear, and weak negative association for rare carnivores. The random intercept standard deviations, a measure of site-level heterogeneity in detections, were significant for all species, with the highest for marmot (SD = 2.16, 95% CI: [1.25- 3.53]) and lowest for rare carnivores (SD = 0.68, 95% CI: [0.03-1.89]; Table S6). All models converged successfully (Rhat <1.10; Table S7; Gelman & Rubin, 1992; Vehtari et al., 2021) and demonstrated good model fit, as indicated by Bayesian p-values near 0.5 and well-aligned posterior predictive checks (Table S8; Figure S8;S9; Gelman et al., 1996).

**Figure 3.**
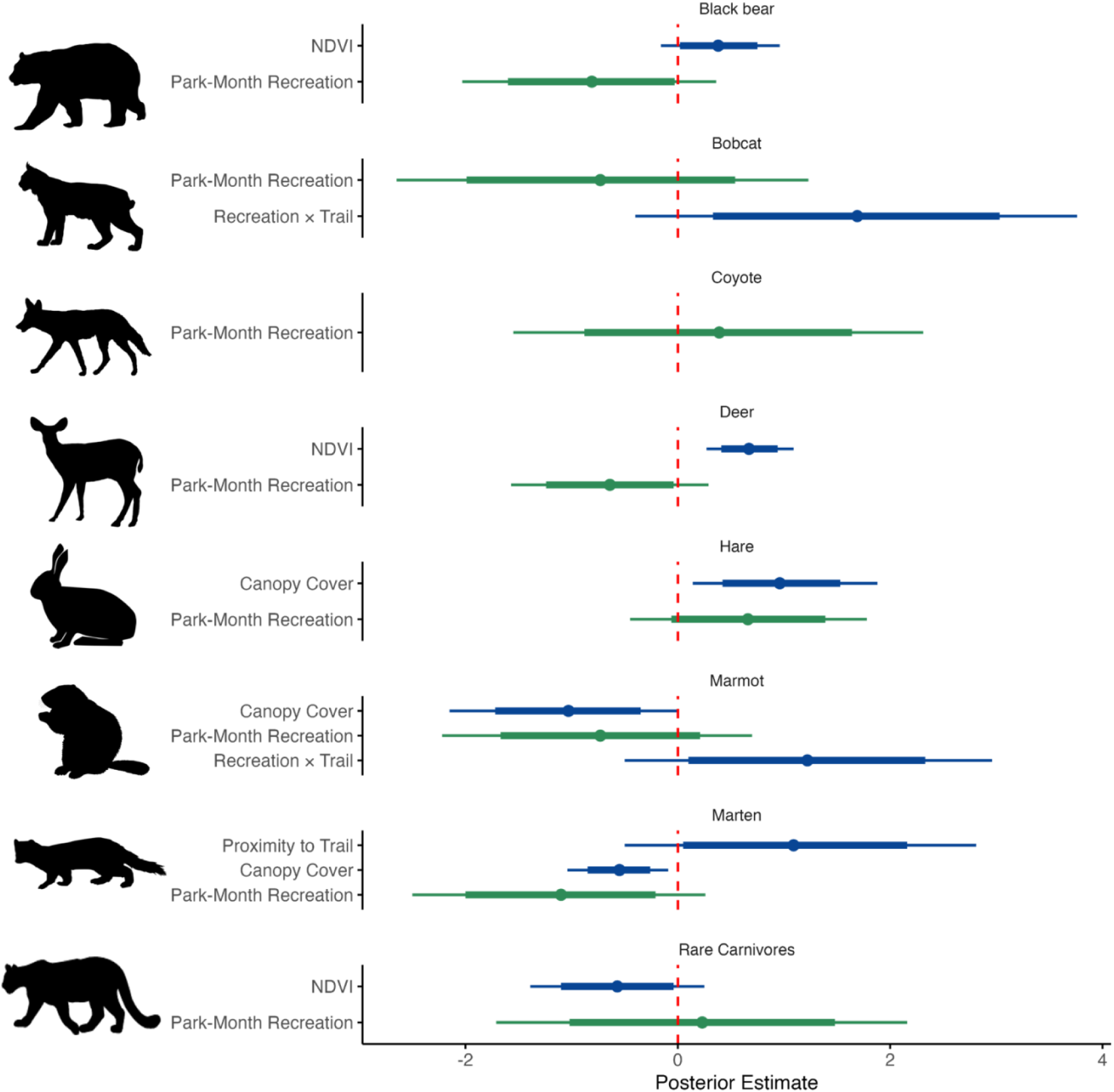
Posterior estimates (x-axis) of effects of Park-Month recreation (green) and other predictor variables for which there was evidence of an association with focal species habitat use (i.e., 80% or 95% credible intervals not overlapping 0; blue) based on Bayesian GLMMs of detections from camera trap surveys in Garibaldi and Joffre Lakes parks. Recreation x Trail represents the interaction term Recreation Park Month x Proximity to Trail (posterior estimates for main effects listed in Table S7). The points represent the mean of the posterior distribution, thick lines represent 95% credible intervals, and the thin lines represent the 80% credible intervals. Parameter estimates for all predictor variables in each species model are listed in Table S7.

### 3.3 Community-level responses to PA closure

In Joffre Lakes, estimated species richness and evenness were not significantly different between years (Figure 4; Table S5). Diversity also remained relatively stable between consecutive years but was significantly greater in 2020 than 2022 (mean difference = −0.38, 95% CI: [−0.59, −0.17], p = 0.02). In Garibaldi, richness decreased significantly from 2020 to 2021 (mean difference = 2.03, 95% CI: [0.89, 3.13], p = 0.02). Diversity and evenness did not change significantly over time. In 2020, species richness (mean difference= −5.04, 95% CI: [−8.36, - 1.69], p = 0.04) and diversity (mean difference = −0.33, 95% CI: [−0.50, −0.16], p = 0.02) were higher in Joffre Lakes than Garibaldi, but did not differ significantly between the parks in the other periods. Evenness stayed relatively consistent between and within the parks, suggesting that the differences in diversity mostly reflected shifts in richness, not evenness (Table S5).

**Figure 4.**
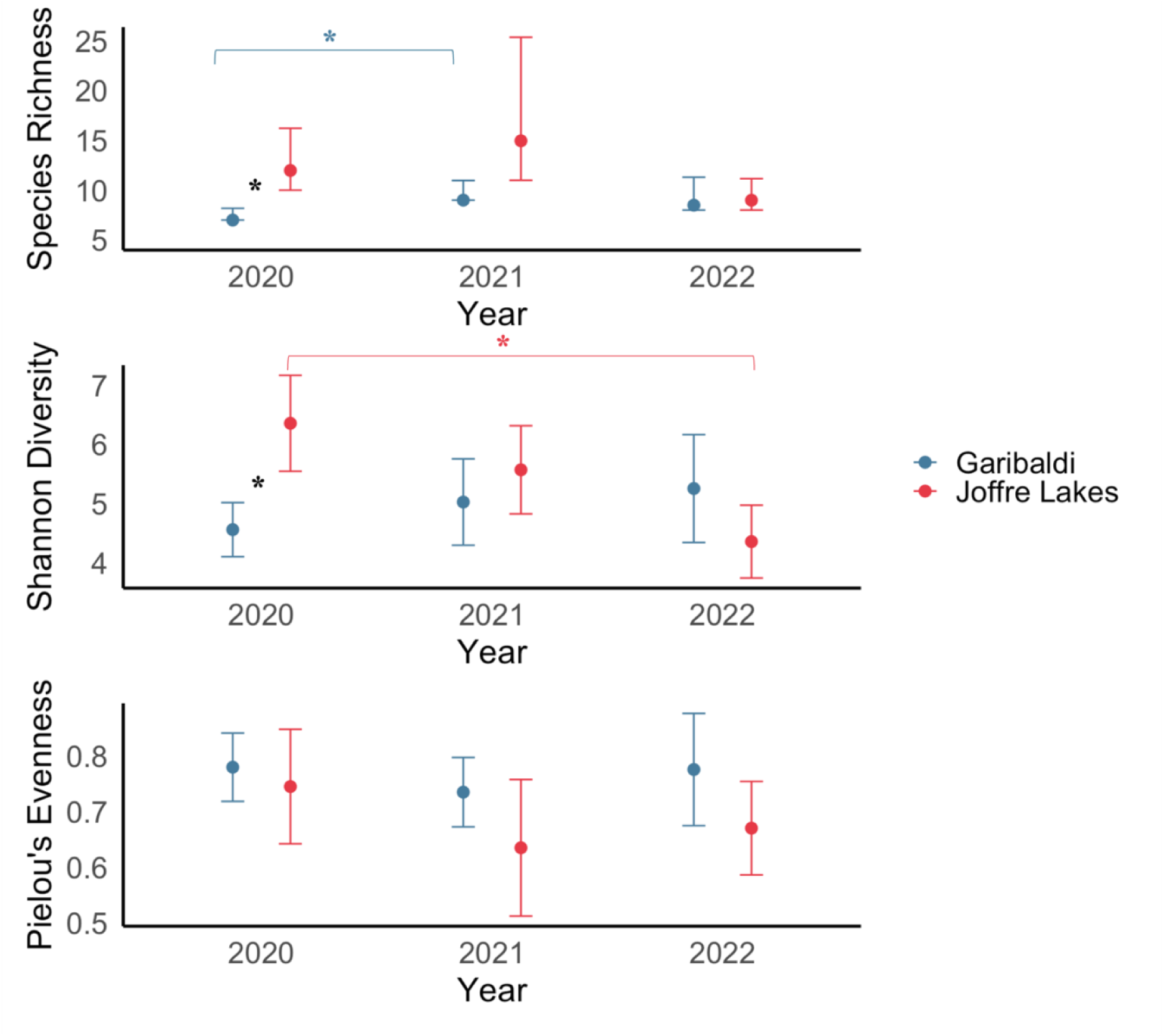
Diversity estimates (points) and confidence intervals (bars) by PA and year (x-axis). Species richness and diversity were estimated using *iNEXT* species accumulation curves for a 90-day period in each year (Figure S10). Evenness was calculated using the species richness and diversity estimates, with confidence intervals propagated from first-order Taylor expansion. Asterisks mark statistically significant differences in diversity metrics, determined by bootstrap hypothesis tests.

## 4. Discussion

### 4.1 Summary of key findings

Our findings revealed mixed evidence for mammal community responses to increasing recreation. We used a quasi-experimental design to leverage dramatic spatial and temporal variation in recreation intensity following COVID-19 closures in two PAs with popular hiking trails. We found evidence that species richness and diversity were highest when and where recreation was lowest: in the closed Joffre Lakes. However, there were inconsistent patterns of community-level changes following the re-opening of Joffre Lakes, with diversity significantly lower in the second year after re-opening, but no significant reductions in richness. Community evenness remained relatively consistent through the study, indicating little change in relative species abundances.

Consistent with this community-level result, none of the more common species in our study exhibited strong spatiotemporal associations with recreation. We identified weak evidence of a negative association between monthly recreation intensity and detections for black bear, marten and deer, and a positive interaction between recreation and proximity to trail for detections of marmot and bobcat. Nevertheless, many of the 15 total species detected were rare in this system, especially those predicted to be highly sensitive (Table 1; S1; S2; e.g., wolves and grizzly bears). The seven more common species for which we could model spatial associations may have been detected more frequently due to their greater tolerance for recreation. These results suggest that behavioural responses to recreation by more common species, such as their habitat use, may not reflect broader changes in community structure that are driven by the responses of rarer and more sensitive species. Our findings provide valuable insights into the impact of recreation on mammal diversity and how future studies should approach assessing both community- and species-level responses to recreation.

### 4.2 Drivers of wildlife habitat use

We found limited support for our prediction that larger-bodied, longer-lived, and wider-ranging species would be more sensitive to recreation. We did not observe an association between the aggregated rare carnivore group and recreation but did observe weak evidence of a negative association between high recreation months and black bear habitat use. In contrast, smaller carnivores exhibited mixed responses. Martens were detected less during high recreation months but were generally detected more near trails regardless of recreation intensity, suggesting a preference for linear features as travel corridors, but an avoidance of busy recreation months. Conversely, bobcats were detected more frequently near or on trails during high-recreation months (Figure S7A). Among the prey species modelled, deer displayed weak evidence of a negative association with high recreation months, and marmots displayed weak evidence of a positive spatial response, using trails more during high-recreation months (Figure S7B).

Despite weak evidence of human avoidance by black bears, we found very limited support for a human shield effect, suggested only in spatial responses by a single subordinate predator (bobcat) and prey species (marmot), which is consistent with other recent studies showing little support for this hypothesis (e.g., Granados et al., 2023; Green et al., 2023). However, it is important to note that other common predators were not detected sufficiently to be modelled (e.g. cougars; Forrester & Wittmer, 2024; Procko et al., 2022) and we could not account for potential confounders such as human behaviour (e.g., food attractants; Gaynor et al., 2025). Both of the smaller, shorter-lived, and smaller-ranging prey species, marmot and hare, had stronger associations with canopy cover than recreation, indicating that structural features and concealment from predators may be more important determinants of habitat use. While hares favoured areas with greater cover (Litvaitis, 1991; Zimova et al., 2014), possibly for greater concealment from their many predators, marmots preferred sites with lower canopy cover. We posit this is unlikely to reflect an active preference for open habitats, but rather their use of alpine environments where the canopy is naturally sparse, and marmots instead use burrows for predator refuge (Blumstein, 2020; Holmes, 1984). In both cases, these structural features offer consistent protection from predation, unlike the temporally variable protection that would be afforded by a human shield effect. In the case of deer, as a larger-bodied, wider-ranging, and longer-lived prey species (Table S1; S2), they are more likely to come into conflict with humans, which may explain why they were detected less during high recreation months altogether, unlike marmots (Visscher et al., 2023; Westekemper et al., 2018). Further, deer may actively avoid humans to protect their young, given their substantial investment in parental care (Faull et al., 2023). Although functional traits and interspecific interactions often influence sensitivity to human disturbance, the consistently weak or neutral responses across species suggest these may not be strong predictors in this context. Notably, our species-specific findings are not consistent with others reported in similar studies conducted in PAs during the pandemic. For example, mule deer were often found to respond positively to high levels of recreation post-closure (Anderson et al., 2023; Fennell et al., 2023; Procko et al., 2022). This highlights the context-dependent nature of wildlife responses to recreation, making generalizations across PAs challenging and an important subject for continued inquiry (Burton et al., 2024; Granados et al., 2023; Marion et al., 2020).

While top predators play a crucial role in regulating community dynamics and are generally expected to be more sensitive to human disturbance (Baker & Leberg, 2018; Ripple et al., 2014), we did not find conclusive evidence for this sensitivity in our study. Since apex predators naturally occur at low densities, detection rates for most of these species were very low (Table S1), necessitating our aggregate rare carnivore model. Future studies may benefit from longer-term and/or more targeted sampling (e.g., using species-specific baits or lures; Lamb et al., 2016; Long et al., 2024), larger CT arrays (Kays, et al., 2020), or multi-array syntheses (Burton et al., 2024; Granados et al., 2023) to increase detection rates of rare carnivores, thereby allowing for robust species-specific modelling,. The low detection rates of these species could also result from insufficient sampling effort, or it may imply that these sensitive species have already begun to be filtered out of busy PAs (Burton et al., 2024; Kays, Arbogast, Baker-Whatton, et al., 2020). Such filtering is further supported by the fact that some of these species (i.e. grey wolf, Canada lynx and grizzly bear) were only detected in Joffre Lakes, which had a longer closure period and consistently lower recreation intensity than Garibaldi (Figure 2).

Additionally, wolves, lynx and grizzly bears were only detected at off-trail CT sites (408, 261, 239 meters from the nearest trail, respectively) and in densely forested areas (i.e., greater concealment cover and sound buffer from human activity; Fricke, 1984; Gaynor & Green, 2024; Zeller et al., 2024) as indicated by high canopy cover (80, 85, 86 percent, respectively). As a much larger PA, Garibaldi offers low disturbance areas outside of the area sampled in our study, whereas sensitive species in Joffre Lakes may be forced to adjust behaviour temporally in order to co-exist with recreationists due to lower availability of alternate habitat (Gill et al., 2001). We suspect that more remote and forested areas of these highly recreated landscapes may offer refuge to these sensitive species, implying the need to maintain low-disturbance areas in popular PAs.

The lack of strong evidence of a response for many species that we expected to avoid times and areas of high recreation is not inconsistent with other behavioural adjustments in response to recreation, such as diel activity partitioning (Anderson et al., 2023; Naidoo & Burton, 2020; Procko et al., 2023), or longer-term adjustments like habituation (Bejder et al., 2009; Berger & Cassidy, 2024; Golden Beam et al., 2023). When wildlife perceive human-dominated areas (e.g., busy trails) as risky, they often adjust their use of the landscape to avoid humans (Frid & Dill, 2002; Gaynor, Hayes, et al., 2025; Palmer et al., 2023). However, this avoidance may come at a cost, limiting access to high-quality resources (Smith et al., 2017; Suraci et al., 2019; Ydenberg & Dill, 1986). Conversely, repeated benign interactions with humans (e.g. during low-impact activities like hiking) can reduce the perceived risk, fostering tolerance and allowing species to prioritize resources over their perceived fear of humans (Blumstein, 2016; Čapkun-Huot et al., 2024; Samia et al., 2015). Decades of high recreation in both PAs before COVID-19 (Kohlhardt et al., 2018; Ministry of Environment and Climate Change Strategy, 2021; Smith et al., 2022) may have weakened the responses of otherwise sensitive species to human disturbance. Habituation can mitigate some of the negative effects of recreation on wildlife but can also lead to unintended consequences such as increased human-wildlife conflict (Barrett et al., 2019; Honda et al., 2014, 2019; Uchida et al., 2024; Wheat & Wilmers, 2016). In order to garner empirical evidence on whether some species habituate after long-term exposure to recreation, long-term monitoring is needed.

### 4.3 Community shifts in response to PA reopening

The community models provided some support for our prediction that richness would be greater under lower recreation. Richness and diversity were indeed higher in Joffre Lakes during the 2020 closure, relative to the open Garibaldi. However, contrary to our prediction, richness did not decrease with increasing recreation in Joffre Lakes in the 2 years following re-opening. Recreation was less variable over time in Garibaldi, but the period with the lowest recreation intensity (2021) had greater species richness than the previous period (2020; Figure 2; 5). In contrast, we found no support for the hypothesized shift in evenness in response to recreation within the two years of this study. This discrepancy may reflect the high proportion of rare species in the community, whose presence or absence influences richness but has a limited impact on relative abundance distribution. The consistency in evenness suggests that the more common or dominant species maintained similar relative abundances across years. This highlights the complexity of interpreting community-level patterns over short timescales, especially in systems with many rare species.

The lack of a significant change in richness over time in Joffre Lakes, despite variation in recreation intensity, may reflect two key considerations for detecting community shifts. First, Chao estimators rely heavily on the frequency of singleton and doubleton detections to extrapolate undetected species and estimate total richness. This can introduce high uncertainty when many rare species are detected (e.g., Joffre Lakes in 2021; Figure 4; Chao et al., 2014), making it difficult to distinguish true community changes from sampling variance. Second, it is possible that community shifts happen more slowly due to a lag in response, similar to delayed effects of disturbance observed in other contexts (e.g., forest fragmentation; Semper-Pascual et al., 2021). For instance, species richness in 2021 may have remained similar to that in 2020 because the community had not yet fully responded to the sudden increase in human presence, considering this sampling period began the day the PA reopened (Chen et al., 2023; Essl et al., 2015b; Wilson et al., 2020). Similarly, recreation levels were lowest in 2020 during COVID-19 closures, but this followed many years of high recreation that may have had lasting effects on species occurrence patterns (Ministry of Environment and Climate Change Strategy, 2021). The fact that we found diversity in Joffre Lakes in 2020 to be higher than in 2022, but not than in 2021, could reflect a modest, lagged increase in species richness in that first year after reopening (though the effect was not statistically significant). This mismatch of time scales may be particularly relevant for longer-lived species, which are likely to be influenced by the cumulative effects of past and current recreation intensity (Jiménez-Franco et al., 2022; McKelvey, 2015). For instance, the two rarest species in our survey, Canada lynx and grizzly bear, which we hypothesized to be sensitive to recreation (Table S1; S2), were only detected in 2021 in Joffre Lakes, just months after the PA reopened. Comparatively, shorter-lived species may respond to disturbances more quickly, as their shorter generation times allow population-level changes to manifest much faster (White et al., 2022 fr sp (Essl et al., 2015a).

### 4.4 Study limitations and constraints

It is often challenging to balance robust research design with practical management constraints, and as such, our quasi-experimental design was not perfect. For instance, some members of the public still used the PAs during official closures, and Joffre Lakes reopened mid-summer in 2021, blurring the lines between open and closed in the context of our seasonal sampling periods. Nonetheless, experimental approaches remain an important means of seeking stronger, causal inference, and thus we call for more collaboration between researchers and PA managers to implement effective adaptive management (Naidoo et al., 2025; Newcomb et al., 2021).

## 5. Conclusion

As demand for outdoor recreation continues to expand globally, understanding its impacts on wildlife is essential for PA management. The COVID-19 closures in PAs served as a novel opportunity to examine wildlife responses to recreation. We found evidence for a decline in species richness in an open PA compared to a closed PA, which may reflect filtering of sensitive species. Yet, habitat use among common species was mostly weakly or neutrally associated with recreation. This pattern is consistent with the absence of any significant change in community evenness, suggesting that community-level responses are likely driven by rare and sensitive species, which are inherently difficult to survey. We know that PAs are critical for protecting biodiversity (Chen et al., 2022). In our study system, most species did not strongly avoid hikers, which implies there is scope for coexistence between low-impact recreation and conservation in PAs. However, rigorous, long-term monitoring is essential to fully understand these dynamics while ensuring that rare species are adequately sampled and effectively protected. As recreation continues to grow in PAs, ecologists and managers must prioritize such monitoring to support adaptive management and informed decision-making.

## TARGET AUDIENCE

recreation ecologists, wildlife biologists, and protected area managers seeking to understand the population- and community-level outcomes of increasing recreation in protected areas, with the goal of improving monitoring efforts and adaptive management.

## AUTHOR CONTRIBUTIONS

**Alexandra Dimitriou**: Conceptualization, Data curation, Formal analysis, Methodology, Writing – Original Draft. **Sarah Benson-Amram**: Conceptualization, Writing – Review & Editing. **Kaitlyn M. Gaynor**: Conceptualization, Writing – Review & Editing. **A. Cole Burton**: Conceptualization, Funding acquisition, Methodology, Project administration, Supervision, Investigation, Writing – Review & Editing.

## ACKNOWLEDGMENTS

We are grateful to have carried out this study on the traditional territories of the Squamish, Lil’wat and N’Quatqua First Nations, who have inhabited and stewarded these lands since time immemorial. We thank members of the UBC Wildlife Coexistence Lab for their efforts in study design, camera trap deployment, data collection, and data processing, including A. Granados, C. Beirne, R. Cameron, C. Chen, J. Clarke, A. Constantinou, M. Fennell, I. Francis, Z. Konanz, M. Procko, C. Sun, and K. Tjaden-McClement. Additionally, we are grateful for feedback on methodology and early versions of the manuscript from members of the Wildlife Coexistence Lab. We also thank the BC Parks team for support, including M. Percy, J. Quayle, J. Hirner, K. Wagner, D. Whiteside and Z. Tulcsik. This research was funded by the BC Ministry of Environment and Parks, and the Natural Sciences and Engineering Council of Canada (Canada Research Chair and Discovery Grant RGPIN-2018-03958 to ACB). AD was supported by a British Columbia Graduate Scholarship (Government of British Columbia), Canada Graduate Scholarship (Natural Sciences and Engineering Council of Canada), and Faculty of Forestry Graduate Award (University of British Columbia).

## ETHICS STATEMENT

This research was conducted under protocol # A25-0104 from the University of British Columbia’s Animal Care Committee and under protocol # H21-01424 from the University of British Columbia’s Behavioral Research Ethics Board.

## References

Anderson, A. K., Waller, J. S., & Thornton, D. H. (2023). Partial COVID-19 closure of a national park reveals negative influence of low-impact recreation on wildlife spatiotemporal ecology. Scientific Reports, 13(1), Article 1. 10.1038/s41598-023-27670-9

Arlettaz, R., Patthey, P., Baltic, M., Leu, T., Schaub, M., Palme, R., & Jenni-Eiermann, S. (2007). Spreading free-riding snow sports represent a novel serious threat for wildlife. Proceedings of the Royal Society B: Biological Sciences, 274(1614), 1219–1224. 10.1098/rspb.2006.0434

Baker, A. D., & Leberg, P. L. (2018). Impacts of human recreation on carnivores in protected areas. PLOS ONE, 13(4), e0195436. 10.1371/journal.pone.0195436

Baluja, T. (2016, September 13). Is social media ruining hiking in B.C.? CBC News. https://www.cbc.ca/news/canada/british-columbia/social-media-hiking-1.3755738

Barrett, L. P., Stanton, L. A., & Benson-Amram, S. (2019). The cognition of ‘nuisance’ species. Animal Behaviour, 147, 167–177. 10.1016/j.anbehav.2018.05.005

Bates, A. E., Primack, R. B., Moraga, P., & Duarte, C. M. (2020). COVID-19 pandemic and associated lockdown as a “Global Human Confinement Experiment” to investigate biodiversity conservation. Biological Conservation, 248, 108665. 10.1016/j.biocon.2020.108665

BC Parks. (n.d.). BC Parks Day-Use Pass Statistic Summary Report 2020-2023. https://nrs.objectstore.gov.bc.ca/kuwyyf/bcparks_day_use_pass_statistics_report_2020_2023_746adb4c3f.pdf

Beery, S., Morris, D., Yang, S., Simon, M., Norouzzadeh, A., & Joshi, N. (2019). Efficient Pipeline for Automating Species ID in new Camera Trap Projects. Biodiversity Information Science and Standards, 3, e37222. 10.3897/biss.3.37222

Beery, T., Olsson, M. R., & Vitestam, M. (2021). Covid-19 and outdoor recreation management: Increased participation, connection to nature, and a look to climate adaptation. Journal of Outdoor Recreation and Tourism, 36, 100457. 10.1016/j.jort.2021.100457

Bejder, L., Samuels, A., Whitehead, H., Finn, H., & Allen, S. (2009). Impact assessment research: Use and misuse of habituation, sensitisation and tolerance in describing wildlife responses to anthropogenic stimuli. Marine Ecology Progress Series, 395, 177–185. 10.3354/meps07979

Benjamini, Y., & Hochberg, Y. (1995). Controlling the False Discovery Rate: A Practical and Powerful Approach to Multiple Testing. Journal of the Royal Statistical Society: Series B (Methodological), 57(1), 289–300. 10.1111/j.2517-6161.1995.tb02031.x

Berger, J. (2007). Fear, human shields and the redistribution of prey and predators in protected areas. Biology Letters, 3(6), 620–623. 10.1098/rsbl.2007.0415

Berger, J., & Cassidy, K. A. (2024). Play is a privilege in both humans and animals: How our recreation influences wildlife. The Journal of Wildlife Management, 88(8), e22664. 10.1002/jwmg.22664

Bergman, J. N., Buxton, R. T., Lin, H.-Y., Lenda, M., Attinello, K., Hajdasz, A. C., Rivest, S. A., Tran Nguyen, T., Cooke, S. J., & Bennett, J. R. (2022). Evaluating the benefits and risks of social media for wildlife conservation. FACETS, 7, 360–397. 10.1139/facets-2021-0112

Berkes, F., Colding, J., & Folke, C. (2008). Navigating Social-Ecological Systems: Building Resilience for Complexity and Change. Cambridge University Press.

Blumstein, D. T. (2016). Habituation and sensitization: New thoughts about old ideas. Animal Behaviour, 120, 255–262. 10.1016/j.anbehav.2016.05.012

Blumstein, D. T. (2020). The Nature of Fear: Survival Lessons from the Wild. Harvard University Press.

Bro-Jørgensen, J., Franks, D. W., & Meise, K. (2019). Linking behaviour to dynamics of populations and communities: Application of novel approaches in behavioural ecology to conservation. Philosophical Transactions of the Royal Society B: Biological Sciences, 374(1781), 20190008. 10.1098/rstb.2019.0008

Bürkner, P.-C. (2017). brms: An R Package for Bayesian Multilevel Models Using Stan. Journal of Statistical Software, 80, 1–28. 10.18637/jss.v080.i01

Burton, A. C., Beirne, C., Gaynor, K. M., Sun, C., Granados, A., Allen, M. L., Alston, J. M., Alvarenga, G. C., Calderón, F. S. Á., Amir, Z., Anhalt-Depies, C., Appel, C., Arroyo-Arce, S., Balme, G., Bar-Massada, A., Barcelos, D., Barr, E., Barthelmess, E. L., Baruzzi, C., … Kays, R. (2024). Mammal responses to global changes in human activity vary by trophic group and landscape. Nature Ecology & Evolution, 1–12. 10.1038/s41559-024-02363-2

Burton, A. C., Neilson, E., Moreira, D., Ladle, A., Steenweg, R., Fisher, J. T., Bayne, E., & Boutin, S. (2015). REVIEW: Wildlife camera trapping: a review and recommendations for linking surveys to ecological processes. Journal of Applied Ecology, 52(3), 675–685. 10.1111/1365-2664.12432

Bushell, R., & McCool, S. (2007). Tourism and protected areas: Benefits beyond boundaries. The Vth IUCN World Parks Congress. https://scholar.google.com/scholar?hl=en&as_sdt=0%2C5&q=bushell+2007+tool+for+conservation&btnG=&oq=bushell+2007

Butchart, S. H. M., Walpole, M., Collen, B., van Strien, A., Scharlemann, J. P. W., Almond, R. E. A., Baillie, J. E. M., Bomhard, B., Brown, C., Bruno, J., Carpenter, K. E., Carr, G. M., Chanson, J., Chenery, A. M., Csirke, J., Davidson, N. C., Dentener, F., Foster, M., Galli, A., … Watson, R. (2010). Global Biodiversity: Indicators of Recent Declines. Science, 328(5982), 1164–1168. 10.1126/science.1187512

Camp, M. J., Rachlow, J. L., Woods, B. A., Johnson, T. R., & Shipley, L. A. (2013). Examining functional components of cover: The relationship between concealment and visibility in shrub-steppe habitat. Ecosphere, 4(2), art19. 10.1890/ES12-00114.1

Čapkun-Huot, C., Blumstein, D. T., Garant, D., Sol, D., & Réale, D. (2024). Toward a unified framework for studying behavioural tolerance. Trends in Ecology & Evolution, 39(5), 446–455. 10.1016/j.tree.2023.12.006

Chao, A., Gotelli, N. J., Hsieh, T. C., Sander, E. L., Ma, K. H., Colwell, R. K., & Ellison, A. M. (2014). Rarefaction and extrapolation with Hill numbers: A framework for sampling and estimation in species diversity studies. Ecological Monographs, 84(1), 45–67. 10.1890/13-0133.1

Chen, C., Brodie, J. F., Kays, R., Davies, T. J., Liu, R., Fisher, J. T., Ahumada, J., McShea, W., Sheil, D., Agwanda, B., Andrianarisoa, M. H., Appleton, R. D., Bitariho, R., Espinosa, S., Grigione, M. M., Helgen, K. M., Hubbard, A., Hurtado, C. M., Jansen, P. A., … Burton, A. C. (2022). Global camera trap synthesis highlights the importance of protected areas in maintaining mammal diversity. Conservation Letters, 15(2), e12865. 10.1111/conl.12865

Chen, X., Wang, Q., Cui, B., Chen, G., Xie, T., & Yang, W. (2023). Ecological time lags in biodiversity response to habitat changes. Journal of Environmental Management, 346, 118965. 10.1016/j.jenvman.2023.118965

Comer, C. E., Bolen, E. G., & Robinson, W. L. (2025). Wildlife Ecology and Management: Sixth Edition. Waveland Press.

Creel, S., Fox, J. E., Hardy, A., Sands, J., Garrott, B., & Peterson, R. O. (2002). Snowmobile Activity and Glucocorticoid Stress Responses in Wolves and Elk. Conservation Biology, 16(3), 809–814. 10.1046/j.1523-1739.2002.00554.x

Crooks, K. R. (2002). Relative Sensitivities of Mammalian Carnivores to Habitat Fragmentation. Conservation Biology, 16(2), 488–502. 10.1046/j.1523-1739.2002.00386.x

DeWan, A. A., & Zipkin, E. F. (2010). An Integrated Sampling and Analysis Approach for Improved Biodiversity Monitoring. Environmental Management, 45(5), 1223–1230. 10.1007/s00267-010-9457-7

Dickie, M., McNay, S. R., Sutherland, G. D., Cody, M., & Avgar, T. (2020). Corridors or risk? Movement along, and use of, linear features varies predictably among large mammal predator and prey species. Journal of Animal Ecology, 89(2), 623–634. 10.1111/1365-2656.13130

Duffy, J. E. (2009). Why biodiversity is important to the functioning of real-world ecosystems. Frontiers in Ecology and the Environment, 7(8), 437–444. 10.1890/070195

Durbin, J., & Watson, G. S. (1950). Testing for Serial Correlation in Least Squares Regression: I. Biometrika, 37(3/4), 409–428. 10.2307/2332391

Efron, B., & Tibshirani, R. J. (1994). An Introduction to the Bootstrap. Chapman and Hall/CRC. 10.1201/9780429246593

Essl, F., Dullinger, S., Rabitsch, W., Hulme, P. E., Pyšek, P., Wilson, J. R. U., & Richardson, D. M. (2015a). Delayed biodiversity change: No time to waste. Trends in Ecology & Evolution, 30(7), 375–378. 10.1016/j.tree.2015.05.002

Essl, F., Dullinger, S., Rabitsch, W., Hulme, P. E., Pyšek, P., Wilson, J. R. U., & Richardson, D. M. (2015b). Historical legacies accumulate to shape future biodiversity in an era of rapid global change. Diversity and Distributions, 21(5), 534–547. 10.1111/ddi.12312

Faull, J., Conteddu, K., Griffin, L. L., Amin, B., Smith, A. F., Haigh, A., & Ciuti, S. (2023). *Does fear of humans predict anti-predator strategies in an ungulate hider species during fawning?* (p. 2023.08.16.553188). bioRxiv. 10.1101/2023.08.16.553188

Fennell, M., Beirne, C., & Burton, A. C. (2022). Use of object detection in camera trap image identification: Assessing a method to rapidly and accurately classify human and animal detections for research and application in recreation ecology. Global Ecology and Conservation, 35, e02104. 10.1016/j.gecco.2022.e02104

Fennell, M., Ford, A., Martin, T., & Burton, C. (2023). Assessing the impacts of recreation on habitat use by mammals in an isolated alpine protected are. 10.22541/au.168673637.72490133/v1

Fennell, M. J. E., Ford, A. T., Martin, T. G., & Burton, A. C. (2023). Assessing the impacts of recreation on the spatial and temporal activity of mammals in an isolated alpine protected area. Ecology and Evolution, 13(11), e10733. 10.1002/ece3.10733

Finney, S. K., Pearce-Higgins, J. W., & Yalden, D. W. (2005). The effect of recreational disturbance on an upland breeding bird, the golden plover *Pluvialis apricaria*. Biological Conservation, 121(1), 53–63. 10.1016/j.biocon.2004.04.009

Fitzpatrick, R. S., Kissel, M. C., Wuensch, M. A., Michael, T. C., & Ward, D. (2024). Anthropogenic linear features exhibit greater mammal activity relative to surrounding game trails in a woody savannah. African Journal of Ecology, 62(3), e13305. 10.1111/aje.13305

Forrester, T. D., & Wittmer, H. (2024). Mountain Lion Predation on Mule and Black-tailed Deer: A Perspective on The Complex Impacts of An Iconic Predator. 10.25455/wgtn.28136150

Fricke, F. (1984). Sound attenuation in forests. Journal of Sound and Vibration, 92(1), 149–158. 10.1016/0022-460X(84)90380-8

Frid, A., & Dill, L. (2002). Human-caused Disturbance Stimuli as a Form of Predation Risk. Conservation Ecology, 6(1). https://www.jstor.org/stable/26271862

Gaynor, K. M., Brown, J. S., Middleton, A. D., Power, M. E., & Brashares, J. S. (2019). Landscapes of Fear: Spatial Patterns of Risk Perception and Response. Trends in Ecology & Evolution, 34(4), 355–368. 10.1016/j.tree.2019.01.004

Gaynor, K. M., & Green, J. R. (2024). Wildlife ecology: The scary sounds of recreation. Current Biology, 34(15), R736–R738. 10.1016/j.cub.2024.06.046

Gaynor, K. M., Hayes, F. P., Manlove, K., Galloway, N., Benson, J. F., Cherry, M. J., Epps, C. W., Fletcher, R. J., Orrock, J., Smith, J. A., Aiello, C., Belant, J. L., Berger, J., Biel, M., Bright, J., Bump, J., Burchett, M., Butler, C., Carlson, J., … Cross, P. C. (2025). The influence of human presence and footprint on animal space use in US national parks. *Proceedings*. Biological Sciences, 292(2051), 20251013. 10.1098/rspb.2025.1013

Gaynor, K. M., Wooster, E. I. F., Martinig, A. R., Green, J. R., Chhen, A., Cuadros, S., Gill, R., Khanal, G., Love, N., Marcus, R., Mills, C. L., Wrensford, K., Wright, N. S., Mezzini, S., Marley, J., & Noonan, M. J. (2025). The Human Shield Hypothesis: Does Predator Avoidance of Humans Create Refuges for Prey? Ecology Letters, 28(6), e70138. 10.1111/ele.70138

Geiser, F. (2013). Hibernation. Current Biology, 23(5), R188–R193. 10.1016/j.cub.2013.01.062

Geldmann, J., Barnes, M., Coad, L., Craigie, I. D., Hockings, M., & Burgess, N. D. (2013). Effectiveness of terrestrial protected areas in reducing habitat loss and population declines. Biological Conservation, 161, 230–238. 10.1016/j.biocon.2013.02.018

Gelman, A. (2006). Prior distributions for variance parameters in hierarchical models (comment on article by Browne and Draper). Bayesian Analysis, 1(3), 515–534. 10.1214/06-BA117A

Gelman, A., Carlin, J. B., Stern, H. S., Dunson, D. B., Vehtari, A., & Rubin, D. B. (2013). *Bayesian Data Analysis* (3rd ed.). Chapman and Hall/CRC. 10.1201/b16018

Gelman, A., Meng, X.-L., & Stern, H. (1996). Posterior Predictive Assessment of Model Fitness Via Realized Discrepancies. Statistica Sinica, 6(4), 733–760.

Gelman, A., & Rubin, D. B. (1992). Inference from Iterative Simulation Using Multiple Sequences. Statistical Science, 7(4), 457–472. 10.1214/ss/1177011136

Gill, J. A., Norris, K., & Sutherland, W. J. (2001). Why behavioural responses may not reflect the population consequences of human disturbance. Biological Conservation, 97(2), 265–268. 10.1016/S0006-3207(00)00002-1

Golden Beam, E. R., Berger, J., Breck, S. W., Schell, C. J., & Lambert, J. E. (2023). Habituation and tolerance in coyotes (Canis latrans), a flexible urban predator. Wildlife Letters, 1(4), 153–162. 10.1002/wll2.12025

Granados, A., Sun, C., Fisher, J. T., Ladle, A., Dawe, K., Beirne, C., Boyce, M. S., Chow, E., Heim, N., Fennell, M., Klees van Bommel, J., Naidoo, R., Procko, M., Stewart, F. E. C., & Burton, A. C. (2023). Mammalian predator and prey responses to recreation and land use across multiple scales provide limited support for the human shield hypothesis. Ecology and Evolution, 13(9), e10464. 10.1002/ece3.10464

Green, A. M., Young, E., Keller, H., Grace, T., Pendergast, M. E., & Şekercioğlu, Ç. H. (2023). Variation in human diel activity patterns mediates periodic increases in recreational activity on mammal behavioural response: Investigating the presence of a temporal ‘weekend effect.’ Animal Behaviour, 198, 117–129. 10.1016/j.anbehav.2023.02.002

Grimm, N. B., Faeth, S. H., Golubiewski, N. E., Redman, C. L., Wu, J., Bai, X., & Briggs, J. M. (2008). Global Change and the Ecology of Cities. Science, 319(5864), 756–760. 10.1126/science.1150195

Gwinn, D. C., Allen, M. S., Bonvechio, K. I., V. Hoyer, M., & Beesley, L. S. (2016). Evaluating estimators of species richness: The importance of considering statistical error rates. Methods in Ecology and Evolution, 7(3), 294–302. 10.1111/2041-210X.12462

Hadidian, J., Prange, S., Rosatte, R., Riley, S. P. D., & Gehrt, S. D. (2010). Raccoons (Procyon lotor). In S. D. Gehrt, S. P. D. Riley, & B. L. Cypher (Eds.), Urban carnivores: Ecology, conflict, and conservation (Vol. 47, pp. 35–48). Johns Hopkins University Press.

Hall, P. (1987). On the Bootstrap and Continuity Correction. Journal of the Royal Statistical Society: Series B (Methodological), 49(1), 82–89. 10.1111/j.2517-6161.1987.tb01428.x

Hill, J. E., DeVault, T. L., Wang, G., & Belant, J. L. (2020). Anthropogenic mortality in mammals increases with the human footprint. Frontiers in Ecology and the Environment, 18(1), 13–18. 10.1002/fee.2127

Hillebrand, H., Blasius, B., Borer, E. T., Chase, J. M., Downing, J. A., Eriksson, B. K., Filstrup, C. T., Harpole, W. S., Hodapp, D., Larsen, S., Lewandowska, A. M., Seabloom, E. W., Van De Waal, D. B., & Ryabov, A. B. (2018). Biodiversity change is uncoupled from species richness trends: Consequences for conservation and monitoring. Journal of Applied Ecology, 55(1), 169–184. 10.1111/1365-2664.12959

Holmes, W. G. (1984). Predation risk and foraging behavior of the hoary marmot in Alaska. Behavioral Ecology and Sociobiology, 15(4), 293–301. 10.1007/BF00292992

Honda, T., Miyagawa, Y., Kuwata, H., Yamasaki, S., & Iijima, H. (2014). Behavioral Traits of Damage-Causing Sika Deer: Open Land Preference. Mammal Study, 39(1), 27–32. 10.3106/041.039.0105

Honda, T., Yamabata, N., Iijima, H., & Uchida, K. (2019). Sensitization to human decreases human-wildlife conflict: Empirical and simulation study. European Journal of Wildlife Research, 65(5), 71. 10.1007/s10344-019-1309-z

Hsieh, T. C., Ma, K. H., & Chao, A. (2016). iNEXT: An R package for rarefaction and extrapolation of species diversity (Hill numbers). Methods in Ecology and Evolution, 7(12), 1451–1456. 10.1111/2041-210X.12613

IUCN. (2024). The IUCN Red List of Threatened Species. Version 2023-3. https://www.iucnredlist.org. Downloaded on 29 February 2024.

Jiménez-Franco, M. V., Graciá, E., Rodríguez-Caro, R. C., Anadón, J. D., Wiegand, T., Botella, F., & Giménez, A. (2022). Problems seeded in the past: Lagged effects of historical land-use changes can cause an extinction debt in long-lived species due to movement limitation. Landscape Ecology, 37(5), 1331–1346. 10.1007/s10980-021-01388-3

Johnson, C. N., Balmford, A., Brook, B. W., Buettel, J. C., Galetti, M., Guangchun, L., & Wilmshurst, J. M. (2017). Biodiversity losses and conservation responses in the Anthropocene. Science, 356(6335), 270–275. 10.1126/science.aam9317

Jones, K. R., Venter, O., Fuller, R. A., Allan, J. R., Maxwell, S. L., Negret, P. J., & Watson, J. E. M. (2018). One-third of global protected land is under intense human pressure. Science, 360(6390), 788–791. 10.1126/science.aap9565

Jost, L. (2010). The Relation between Evenness and Diversity. Diversity, 2(2), 207–232. 10.3390/d2020207

Kays, R., Arbogast, B. S., Baker-Whatton, M., Beirne, C., Boone, H. M., Bowler, M., Burneo, S. F., Cove, M. V., Ding, P., Espinosa, S., Gonçalves, A. L. S., Hansen, C. P., Jansen, P. A., Kolowski, J. M., Knowles, T. W., Lima, M. G. M., Millspaugh, J., McShea, W. J., Pacifici, K., … Spironello, W. R. (2020). An empirical evaluation of camera trap study design: How many, how long and when? Methods in Ecology and Evolution, 11(6), 700–713. 10.1111/2041-210X.13370

Kays, R., Kranstauber, B., Jansen, P., Carbone, C., Rowcliffe, M., Fountain, T., & Tilak, S. (2009). Camera traps as sensor networks for monitoring animal communities. 2009 IEEE 34th Conference on Local Computer Networks, 811–818. 10.1109/LCN.2009.5355046

Kéry, M., & Royle, J. A. (2020). Applied Hierarchical Modeling in Ecology: Analysis of Distribution, Abundance and Species Richness in R and BUGS: Volume 2: Dynamic and Advanced Models. Academic Press.

Kéry, M., & Schmidt, B. (2008). Imperfect detection and its consequences for monitoring for conservation. Community Ecology, 9(2), 207–216. 10.1556/ComEc.9.2008.2.10

Kohlhardt, R., Honey-Rosés, J., Fernandez Lozada, S., Haider, W., & Stevens, M. (2018). Is this trail too crowded? A choice experiment to evaluate tradeoffs and preferences of park visitors in Garibaldi Park, British Columbia. Journal of Environmental Planning and Management, 61(1), 1–24. 10.1080/09640568.2017.1284047

Kunze, C., Bahlburg, D., Urrutia-Cordero, P., Striebel, M., Kelpsiene, E., Langenheder, S., Donohue, I., & Hillebrand, H. (2025). Partitioning species contributions to ecological stability in disturbed communities. Ecological Monographs, 95(1), e1636. 10.1002/ecm.1636

Lacher, T. E., Jr., Davidson, A. D., Fleming, T. H., Gómez-Ruiz, E. P., McCracken, G. F., Owen-Smith, N., Peres, C. A., & Vander Wall, S. B. (2019). The functional roles of mammals in ecosystems. Journal of Mammalogy, 100(3), 942–964. 10.1093/jmammal/gyy183

Lamb, C. T., Walsh, D. A., & Mowat, G. (2016). Factors influencing detection of grizzly bears at genetic sampling sites. Ursus, 27(1), 31–44. 10.2192/URSUS-D-15-00025.1

Larson, C. L., Reed, S. E., Merenlender, A. M., & Crooks, K. R. (2016). Effects of Recreation on Animals Revealed as Widespread through a Global Systematic Review. PLOS ONE, 11(12), e0167259. 10.1371/journal.pone.0167259

Laundré, J. W., Hernández, L., & Altendorf, K. B. (2001). Wolves, elk, and bison: Reestablishing the “landscape of fear” in Yellowstone National Park, U.S.A. Canadian Journal of Zoology, 79(8), 1401–1409. 10.1139/z01-094

Litvaitis, J. A. (1991). Habitat use by Snowshoe Hares, Lepili americanus, in relation to pelage color. The Canadian Field-Naturalist, 105(2), 275--277. 10.5962/p.358009

Long, R. A., MacKay, P., Sauder, J. D., Sinclair, M., Aubry, K. B., & Raley, C. M. (2024). An overwinter protocol for detecting wolverines and other carnivores at camera traps paired with automated scent dispensers. Ecology and Evolution, 14(5), e11290. 10.1002/ece3.11290

Longshore, K., Lowrey, C., & Thompson, D. B. (2013). Detecting short-term responses to weekend recreation activity: Desert bighorn sheep avoidance of hiking trails. Wildlife Society Bulletin, 37(4), 698–706. 10.1002/wsb.349

Mainini, B., Neuhaus, P., & Ingold, P. (1993). Behaviour of marmots *marmota marmota* under the influence of different hiking activities. Biological Conservation, 64(2), 161–164. 10.1016/0006-3207(93)90653-I

Marion, S., Davies, A., Demšar, U., Irvine, R. J., Stephens, P. A., & Long, J. (2020). A systematic review of methods for studying the impacts of outdoor recreation on terrestrial wildlife. Global Ecology and Conservation, 22, e00917. 10.1016/j.gecco.2020.e00917

Matasci, G., Hermosilla, T., Wulder, M. A., White, J. C., Coops, N. C., Hobart, G. W., Bolton, D. K., Tompalski, P., & Bater, C. W. (2018). Three decades of forest structural dynamics over Canada’s forested ecosystems using Landsat time-series and lidar plots. Remote Sensing of Environment, 216, 697–714. 10.1016/j.rse.2018.07.024

McKelvey, K. S. (2015). The effects of disturbance and succession on wildlife habitat and animal communities [Chapter 11]. In: Morrison, M. L.; Mathewson, H. A., Editors. Wildlife Habitat Conservation: Concepts, Challenges, and Solutions. Baltimore, MD: *Johns Hopkins University Press*. p. 143-156., 143–156.

McKinney, M. L., & Lockwood, J. L. (1999). Biotic homogenization: A few winners replacing many losers in the next mass extinction. Trends in Ecology & Evolution, 14(11), 450–453. 10.1016/S0169-5347(99)01679-1

Ministry of Environment and Climate Change Strategy. (2021, June 15). Joffre Lakes Park Visitor Use Management Strategy. https://nrs.objectstore.gov.bc.ca/kuwyyf/joffre_lakes_park_visitor_use_management_strategy_2021_449f5f7368.pdf

Mitterwallner, V., Peters, A., Edelhoff, H., Mathes, G., Nguyen, H., Peters, W., Heurich, M., & Steinbauer, M. J. (2024). Automated visitor and wildlife monitoring with camera traps and machine learning. Remote Sensing in Ecology and Conservation, 10(2), 236–247. 10.1002/rse2.367

Monteith, K. L., Bleich, V. C., Stephenson, T. R., Pierce, B. M., Conner, M. M., Klaver, R. W., & Bowyer, R. T. (2011). Timing of seasonal migration in mule deer: Effects of climate, plant phenology, and life-history characteristics. Ecosphere, 2(4), art47. 10.1890/ES10-00096.1

Naidoo, R., & Burton, A. C. (2020). Relative effects of recreational activities on a temperate terrestrial wildlife assemblage. Conservation Science and Practice, 2(10), e271. 10.1111/csp2.271

Naidoo, R., Gulati, S., Vercammen, J., & Burton, C. (2025). Estimating causal impacts of human recreation on wildlife in the absence of experimental controls. Conservation Letters. In press.

Naylor, L. M., Wisdom, M. J., & Anthony, R. G. (2009). Behavioral Responses of North American Elk to Recreational Activity. The Journal of Wildlife Management, 73(3), 328–338. 10.2193/2008-102

Neilson, E. W., Avgar, T., Burton, A. C., Broadley, K., & Boutin, S. (2018). Animal movement affects interpretation of occupancy models from camera-trap surveys of unmarked animals. Ecosphere, 9(1), e02092. 10.1002/ecs2.2092

Newbold, T., Hudson, L. N., Contu, S., Hill, S. L. L., Beck, J., Liu, Y., Meyer, C., Phillips, H. R. P., Scharlemann, J. P. W., & Purvis, A. (2018). Widespread winners and narrow-ranged losers: Land use homogenizes biodiversity in local assemblages worldwide. PLOS Biology, 16(12), e2006841. 10.1371/journal.pbio.2006841

Newcomb, T. J., Simonin, P. W., Martinez, F. A., Chadderton, W. L., Bossenbroek, J. M., Cudmore, B., Hoff, M. H., Keller, R. P., Ridenhour, B. D., Rothlisberger, J. D., Rutherford, E. S., Van Egeren, S., & Lodge, D. M. (2021). A Best Practices Case Study for Scientific Collaboration between Researchers and Managers. Fisheries, 46(3), 131–138. 10.1002/fsh.10536

Newsome, T. M., & Van Eeden, L. M. (2017). The Effects of Food Waste on Wildlife and Humans. Sustainability, 9(7), Article 7. 10.3390/su9071269

Nickel, B. A., Suraci, J. P., Allen, M. L., & Wilmers, C. C. (2020). Human presence and human footprint have non-equivalent effects on wildlife spatiotemporal habitat use. Biological Conservation, 241, 108383. 10.1016/j.biocon.2019.108383

Ordiz, A., Aronsson, M., Persson, J., Støen, O.-G., Swenson, J. E., & Kindberg, J. (2021). Effects of Human Disturbance on Terrestrial Apex Predators. Diversity, 13(2), Article 2. 10.3390/d13020068

Pacifici, M., Di Marco, M., & Watson, J. E. M. (2020). Protected areas are now the last strongholds for many imperiled mammal species. Conservation Letters, 13(6), e12748. 10.1111/conl.12748

Palmer, M. S., Gaynor, K. M., Abraham, J. O., & Pringle, R. M. (2023). The role of humans in dynamic landscapes of fear. Trends in Ecology & Evolution, 38(3), 217–218. 10.1016/j.tree.2022.12.007

Pettorelli, N., Ryan, S., Mueller, T., Bunnefeld, N., Jedrzejewsk, B., Lima, M., & Kausrud, K. (2011). The Normalized Difference Vegetation Index (NDVI): Unforeseen successes in animal ecology. Climate Research, 46, 15–27. 10.3354/cr00936

Pettorelli, N., Vik, J. O., Mysterud, A., Gaillard, J.-M., Tucker, C. J., & Stenseth, N. Chr. (2005). Using the satellite-derived NDVI to assess ecological responses to environmental change. Trends in Ecology & Evolution, 20(9), 503–510. 10.1016/j.tree.2005.05.011

Pielou, E. C. (1966). The measurement of diversity in different types of biological collections. Journal of Theoretical Biology, 13, 131–144. 10.1016/0022-5193(66)90013-0

Piñeiro, A., Barja, I., Silván, G., & Illera, J. C. (2012). Effects of tourist pressure and reproduction on physiological stress response in wildcats: Management implications for species conservation. Wildlife Research, 39(6), 532–539. 10.1071/WR10218

Potash, A. D., Conner, L. M., & McCleery, R. A. (2019). Vertical and horizontal vegetation cover synergistically shape prey behaviour. Animal Behaviour, 152, 39–44. 10.1016/j.anbehav.2019.04.007

Procko, M., Naidoo, R., LeMay, V., & Burton, A. C. (2022). Human impacts on mammals in and around a protected area before, during, and after COVID-19 lockdowns. Conservation Science and Practice, 4(7), e12743. 10.1111/csp2.12743

Procko, M., Naidoo, R., LeMay, V., & Burton, A. C. (2023). Human presence and infrastructure impact wildlife nocturnality differently across an assemblage of mammalian species. PLOS ONE, 18(5), e0286131. 10.1371/journal.pone.0286131

Prugh, L. R., Cunningham, C. X., Windell, R. M., Kertson, B. N., Ganz, T. R., Walker, S. L., & Wirsing, A. J. (2023). Fear of large carnivores amplifies human-caused mortality for mesopredators. Science, 380(6646), 754–758. 10.1126/science.adf2472

R Core Team. (2024). R: A Language and Environment for Statistical Computing (Version 4.4.0) [Computer software]. https://www.R-project.org/

Reed, S. E., & Merenlender, A. M. (2008). Quiet, Nonconsumptive Recreation Reduces Protected Area Effectiveness. Conservation Letters, 1(3), 146–154. 10.1111/j.1755-263X.2008.00019.x

Ripple, W. J., Estes, J. A., Beschta, R. L., Wilmers, C. C., Ritchie, E. G., Hebblewhite, M., Berger, J., Elmhagen, B., Letnic, M., Nelson, M. P., Schmitz, O. J., Smith, D. W., Wallach, A. D., & Wirsing, A. J. (2014). Status and Ecological Effects of the World’s Largest Carnivores. Science, 343(6167), 1241484. 10.1126/science.1241484

Ritchie, E. G., & Johnson, C. N. (2009). Predator interactions, mesopredator release and biodiversity conservation. Ecology Letters, 12(9), 982–998. 10.1111/j.1461-0248.2009.01347.x

Roswell, M., Dushoff, J., & Winfree, R. (2021). A conceptual guide to measuring species diversity. Oikos, 130(3), 321–338. 10.1111/oik.07202

Rovero, F., & Zimmerman, F. (2016). Camera Trapping for Wildlife Research.

Rowcliffe, J. M., Field, J., Turvey, S. T., & Carbone, C. (2008). Estimating animal density using camera traps without the need for individual recognition. Journal of Applied Ecology, 45(4), 1228–1236. 10.1111/j.1365-2664.2008.01473.x

Rutz. (n.d.). COVID-19 lockdown allows researchers to quantify the effects of human activity on wildlife | Nature Ecology & Evolution. Retrieved April 17, 2025, from https://www.nature.com/articles/s41559-020-1237-z

Rutz, C. (2022). Studying pauses and pulses in human mobility and their environmental impacts. Nature Reviews Earth & Environment, 3(3), 157–159. 10.1038/s43017-022-00276-x

Salvatori, M., Oberosler, V., Rinaldi, M., Franceschini, A., Truschi, S., Pedrini, P., & Rovero, F. (2023). Crowded mountains: Long-term effects of human outdoor recreation on a community of wild mammals monitored with systematic camera trapping. Ambio, 52(6), 1085–1097. 10.1007/s13280-022-01825-w

Samia, D. S. M., Nakagawa, S., Nomura, F., Rangel, T. F., & Blumstein, D. T. (2015). Increased tolerance to humans among disturbed wildlife. Nature Communications, 6(1), 8877. 10.1038/ncomms9877

Santini, L., González-Suárez, M., Russo, D., Gonzalez-Voyer, A., von Hardenberg, A., & Ancillotto, L. (2019). One strategy does not fit all: Determinants of urban adaptation in mammals. Ecology Letters, 22(2), 365–376. 10.1111/ele.13199

Schulze, K., Knights, K., Coad, L., Geldmann, J., Leverington, F., Eassom, A., Marr, M., Butchart, S. H. M., Hockings, M., & Burgess, N. D. (2018). An assessment of threats to terrestrial protected areas. Conservation Letters, 11(3), e12435. 10.1111/conl.12435

Semper-Pascual, A., Burton, C., Baumann, M., Decarre, J., Gavier-Pizarro, G., Gómez-Valencia, B., Macchi, L., Mastrangelo, M. E., Pötzschner, F., Zelaya, P. V., & Kuemmerle, T. (2021). How do habitat amount and habitat fragmentation drive time-delayed responses of biodiversity to land-use change? Proceedings of the Royal Society B: Biological Sciences, 288(1942), 20202466. 10.1098/rspb.2020.2466

Sévêque, A., Gentle, L. K., López-Bao, J. V., Yarnell, R. W., & Uzal, A. (2020). Human disturbance has contrasting effects on niche partitioning within carnivore communities. Biological Reviews, 95(6), 1689–1705. 10.1111/brv.12635

Silva-Rodríguez, E. A., Cortés, E. I., Vasquez-Ibarra, V., Gálvez, N., Cusack, J., Ohrens, O., Moreira-Arce, D., Farías, A. A., & Infante-Varela, J. (2025). A protocol for error prevention and quality control in camera trap datasets. Journal of Applied Ecology. 10.1111/1365-2664.70010

Smith, J. A., McDaniels, M. E., Peacor, S. D., Bolas, E. C., Cherry, M. J., Dorn, N. J., Feldman, O. K., Kimbro, D. L., Leonhardt, E. K., Peckham, N. E., Sheriff, M. J., & Gaynor, K. M. (2024). Population and community consequences of perceived risk from humans in wildlife. Ecology Letters, 27(6), e14456. 10.1111/ele.14456

Smith, J. A., Suraci, J. P., Clinchy, M., Crawford, A., Roberts, D., Zanette, L. Y., & Wilmers, C. C. (2017). Fear of the human ‘super predator’ reduces feeding time in large carnivores. Proceedings of the Royal Society B: Biological Sciences. 10.1098/rspb.2017.0433

Smith, T., Dan, K., & Bulkan, J. (2022). ‘Loved to Death’: Conflicts between Indigenous food sovereignty, settler recreation, and ontologies of land in the governance of Líl̓wat tmicw. BC Studies: The British Columbian Quarterly, 216, Article 216. 10.14288/bcs.no216.196947

Soria, C. D., Pacifici, M., Di Marco, M., Stephen, S. M., & Rondinini, C. (2021). COMBINE: a coalesced mammal database of intrinsic and extrinsic traits.

Suraci, J. P., Clinchy, M., Zanette, L. Y., & Wilmers, C. C. (2019). Fear of humans as apex predators has landscape-scale impacts from mountain lions to mice. Ecology Letters, 22(10), 1578–1586. 10.1111/ele.13344

Suraci, J. P., Gaynor, K. M., Allen, M. L., Alexander, P., Brashares, J. S., Cendejas-Zarelli, S., Crooks, K., Elbroch, L. M., Forrester, T., Green, A. M., Haight, J., Harris, N. C., Hebblewhite, M., Isbell, F., Johnston, B., Kays, R., Lendrum, P. E., Lewis, J. S., McInturff, A., … Wilmers, C. C. (2021). Disturbance type and species life history predict mammal responses to humans. Global Change Biology, 27(16), 3718–3731. 10.1111/gcb.15650

Tattersall, E. R., Burgar, J. M., Fisher, J. T., & Burton, A. C. (2020). Mammal seismic line use varies with restoration: Applying habitat restoration to species at risk conservation in a working landscape. Biological Conservation, 241, 108295. 10.1016/j.biocon.2019.108295

Thompson, P. R., Paczkowski, J., Whittington, J., & St. Clair, C. C. (2025). Integrating human trail use in montane landscapes reveals larger zones of human influence for wary carnivores. Journal of Applied Ecology, 62(2), 344–359. 10.1111/1365-2664.14837

Tuck, S. L., Phillips, H. R. P., Hintzen, R. E., Scharlemann, J. P. W., Purvis, A., & Hudson, L. N. (2014). MODISTools – downloading and processing MODIS remotely sensed data in R. Ecology and Evolution, 4(24), 4658–4668. 10.1002/ece3.1273

Uchida, K., Blumstein, D. T., & Soga, M. (2024). Managing wildlife tolerance to humans for ecosystem goods and services. Trends in Ecology & Evolution, 39(3), 248–257. 10.1016/j.tree.2023.10.008

Vargas Soto, J. S., Beirne, C., Whitworth, A., Cruz Diaz, J. C., Flatt, E., Pillco-Huarcaya, R., Olson, E. R., Azofeifa, A., Saborío-R, G., Salom-Pérez, R., Espinoza-Muñoz, D., Hay, L., Whittaker, L., Roldán, C., Bedoya-Arrieta, R., Broadbent, E. N., & Molnár, P. K. (2022). Human disturbance and shifts in vertebrate community composition in a biodiversity hotspot. Conservation Biology, 36(2), e13813. 10.1111/cobi.13813

Vehtari, A., Gelman, A., Simpson, D., Carpenter, B., & Bürkner, P.-C. (2021). Rank-Normalization, Folding, and Localization: An Improved R^ for Assessing Convergence of MCMC (with Discussion). Bayesian Analysis, 16(2), 667–718. 10.1214/20-BA1221

Vellend, M. (2017). The Biodiversity Conservation Paradox. American Scientist, 105(2), 94. 10.1511/2017.125.94

Visscher, D. R., Walker, P. D., Flowers, M., Kemna, C., Pattison, J., & Kushnerick, B. (2023). Human impact on deer use is greater than predators and competitors in a multiuse recreation area. Animal Behaviour, 197, 61–69. 10.1016/j.anbehav.2023.01.003

Watson, J. E. M., Dudley, N., Segan, D. B., & Hockings, M. (2014). The performance and potential of protected areas. Nature, 515(7525), 67–73. 10.1038/nature13947

Wearn, O. R., & Glover-Kapfer, P. (2019). Snap happy: Camera traps are an effective sampling tool when compared with alternative methods. Royal Society Open Science, 6(3). 10.1098/rsos.181748

Weetman, G., & Prescott, C. (2001). The Structure, Functioning and Management of Old-growth Cedar-Hemlock-Fir Forests on Vancouver Island, British Columbia. In J. Evans (Ed.), The Forests Handbook*, Volume* 2 (pp. 275–287). Blackwell Science Ltd. 10.1002/9780470757079.ch13

Westekemper, K., Reinecke, H., Signer, J., Meißner, M., Herzog, S., & Balkenhol, N. (2018). Stay on trails – effects of human recreation on the spatiotemporal behavior of red deer Cervus elaphus in a German national park. Wildlife Biology, 2018(1), wlb.00403. 10.2981/wlb.00403

Weststrate, D. K., Chhen, A., Mezzini, S., Safford, K., & Noonan, M. J. (2025). How climate change and population growth will shape attendance and human-wildlife interactions at British Columbia parks. Journal of Sustainable Tourism, 33(2), 318–332. 10.1080/09669582.2024.2331228

Wheat, R. E., & Wilmers, C. C. (2016). Habituation reverses fear-based ecological effects in brown bears (Ursus arctos). Ecosphere, 7(7), e01408. 10.1002/ecs2.1408

White, J. W., Barceló, C., Hastings, A., & Botsford, L. W. (2022). Pulse disturbances in age-structured populations: Life history predicts initial impact and recovery time. Journal of Animal Ecology, 91(12), 2370–2383. 10.1111/1365-2656.13828

Wilson, M. W., Ridlon, A. D., Gaynor, K. M., Gaines, S. D., Stier, A. C., & Halpern, B. S. (2020). Ecological impacts of human-induced animal behaviour change. Ecology Letters, 23(10), 1522–1536. 10.1111/ele.13571

Winter, P. L., Selin, S., Cerveny, L., & Bricker, K. (2020). Outdoor Recreation, Nature-Based Tourism, and Sustainability. Sustainability, 12(1), Article 1. 10.3390/su12010081

Ydenberg, R. C., & Dill, L. M. (1986). The Economics of Fleeing from Predators. In J. S. Rosenblatt, C. Beer, M.-C. Busnel, & P. J. B. Slater (Eds.), Advances in the Study of Behavior (Vol. 16, pp. 229–249). Academic Press. 10.1016/S0065-3454(08)60192-8

Zeller, K. A., Ditmer, M. A., Squires, J. R., Rice, W. L., Wilder, J., DeLong, D., Egan, A., Pennington, N., Wang, C. A., Plucinski, J., & Barber, J. R. (2024). Experimental recreationist noise alters behavior and space use of wildlife. Current Biology, 34(13), 2997–3004.e3. 10.1016/j.cub.2024.05.030

Zimova, M., Mills, L. S., Lukacs, P. M., & Mitchell, M. S. (2014). Snowshoe hares display limited phenotypic plasticity to mismatch in seasonal camouflage. Proceedings of the Royal Society B: Biological Sciences, 281(1782), 20140029. 10.1098/rspb.2014.0029

