## Supporting Information for "The effect of outdoor recreation on mammal habitat use and diversity revealed by COVID-19 closures"

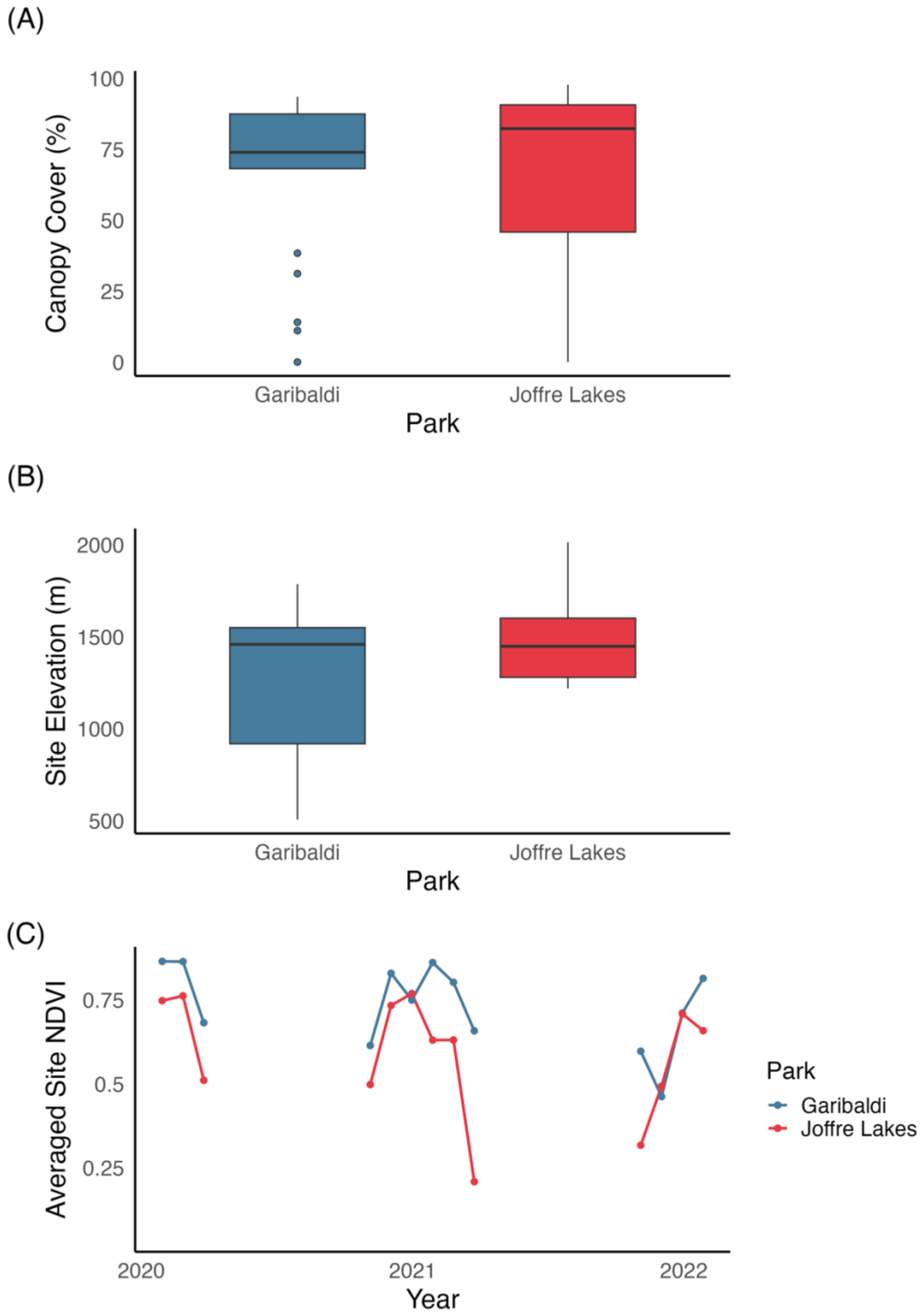

**Figure S1.** Comparison of environmental characteristics between Garibaldi (blue) and Joffre Lakes (red) parks. A) Elevation (in meters) by CT site in each park. B) Canopy cover (percent of vegetation above 2m from NTEMS) by CT site in each park. C) Average Site NDVI by park and month. The contrast in average site NDVI between parks in late 2021 and early 2022 can likely be attributed to earlier snowfall and later snowmelt in the higher elevation sites in Joffre Lakes.

**Table S1.** Summary of species detected in Joffre Lakes and Garibaldi, including number of total detections, which PAs they were detected in in this study (JL = Joffre Lakes, GB = Garibaldi), scientific and common name, taxonomic order, trophic level (1= herbivore, 2= omnivore, 3= carnivore), diet breadth (the number of dietary categories consumed), habitat breadth (number of suitable level 1 IUCN habitats the species uses), generation length in days, and dispersal in kilometers. Row colour is based on trait-based sensitivity scores in table S2 (i.e., darker = more sensitive). Species are ordered by total detections in descending order. Trait data are derived from COMBINE: a coalesced mammal database of intrinsic and extrinsic traits compiled by Soria et al. (2021).

| Species | Common Name | Order | Total Detections | PAs Detected | Adult body mass (g) | Trophic level | Diet breadth | Habitat breadth | Generation length | Dispersal (km) |
|---|---|---|---|---|---|---|---|---|---|---|
| *Lepus americanus* | Snowshoe hare | Lagomorpha | 859 | JL, GB | 1,565.61 | 1 | 1 | 2 | 1,105.95 | 1.36 |
| *Odocoileus hemionus* | Mule deer | Artiodactyla | 730 | JL, GB | 57,000 | 1 | 1 | 9 | 2,717.43 | 9.80 |
| *Ursus americanus* | Black bear | Carnivora | 249 | JL, GB | 132,405 | 2 | 2 | 4 | 4,734.08 | 15.57 |
| *Martes americana* | American marten | Carnivora | 145 | JL, GB | 892 | 2 | 2 | 1 | 2,227.96 | 0.99 |
| *Marmota caligata* | Hoary marmot | Rodentia | 48 | JL, GB | 4,298.52 | 1 | 3 | 2 | 1,906.50 | 2.36 |

| | | | | | | | | | | |
|---|---|---|---|---|---|---|---|---|---|---|
| *Lynx rufus* | Bobcat | Carnivora | 42 | JL, GB | 9,400 | 3 | 1 | 4 | 3,810.03 | 19.49 |
| *Canis latrans* | Coyote | Carnivora | 35 | JL, GB | 11,050 | 3 | 2 | 5 | 2,569.03 | 21.09 |
| *Neotoma cinerea* | Bushy-tailed woodrat | Rodentia | 34 | JL | 285.89 | 1 | 2 | 5 | 1,258.38 | 0.53 |
| *Puma concolor* | Cougar | Carnivora | 15 | JL, GB | 48,000 | 3 | 1 | 5 | 2,693.79 | 43.32 |
| *Erethizon dorsatum* | Porcupine | Rodentia | 10 | JL | 8,009.73 | 1 | 2 | 3 | 3,011.16 | 3.33 |
| *Gulo gulo* | Wolverine | Carnivora | 8 | JL, GB | 12,792.49 | 2 | 4 | 4 | 2,622.30 | 4.31 |
| *Canis lupus* | Grey wolf | Carnivora | 5 | JL | 29,190.76 | 3 | 1 | 7 | 2,883.26 | 33.95 |
| *Procyon lotor* | Raccoon | Carnivora | 4 | GB | 5,075 | 2 | 4 | 1 | 2,703.92 | 2.59 |
| *Lynx canadensis* | Canada lynx | Carnivora | 1 | JL | 9,682.82 | 3 | 2 | 2 | 3,391.96 | 19.77 |
| *Ursus arctos* | Grizzly bear | Carnivora | 1 | JL | 240,500 | 2 | 2 | 5 | 5,976.27 | 21.62 |

**Table S2.** Trait-based sensitivity scores for all mammal species detected based on ecological traits: body mass, generation length, dispersal, trophic level, diet breadth, and habitat breadth (Table 1; Table S1). The following traits: body mass, generation length, dispersal, and trophic level were scored out of 3 points, and niche breadth was scored out of 3 (1.5 for diet and 1.5 for habitat breadth) to allow a maximum total score of 15. Final sensitivity levels were assigned based on total score using the following thresholds: High (≥ 10.5), Moderate–High (7–10.49), Moderate (4.5–6.99), and Low (< 4.5). Note that these scores represent intrinsic ecological traits and do not capture behavioural adaptations that may also influence species' sensitivity to anthropogenic disturbance. They should be interpreted as a baseline measure of sensitivity, considered in conjunction with extrinsic factors like behaviour (e.g. raccoons were scored as moderately sensitive but are well-known synanthropic urban opportunists and are therefore less sensitive to humans; Hadidian et al., 2010).

| **Species** | **Body Mass (0–3)** | **Generation Length (0–3)** | **Dispersal (0–3)** | **Trophic Level (0–3)** | **Diet Breadth (0–1.5)** | **Habitat Breadth (0–1.5)** | **Total Score** | **Sensitivity Level** |
|---|---|---|---|---|---|---|---|---|
| *Puma concolor* | 2 | 1 | 3 | 3 | 1.5 | 0.75 | 11.25 | High |
| *Canis lupus* | 2 | 1 | 3 | 3 | 1.5 | 0.0 | 10.5 | High |
| *Ursus americanus* | 3 | 3 | 2 | 1 | 0.75 | 0.75 | 10.5 | High |
| *Ursus arctos* | 3 | 3 | 2 | 1 | 0.75 | 0.75 | 10.5 | High |
| *Lynx rufus* | 1 | 2 | 2 | 3 | 1.5 | 0.75 | 10.25 | Moderate–High |
| *Lynx canadensis* | 1 | 2 | 2 | 3 | 0.75 | 1.5 | 10.25 | Moderate–High |
| *Canis latrans* | 1 | 1 | 2 | 3 | 0.75 | 0.75 | 8.5 | Moderate–High |
| *Odocoileus hemionus* | 2 | 1 | 1 | 0 | 1.5 | 0.0 | 5.5 | Moderate |
| *Erethizon dorsatum* | 1 | 2 | 1 | 0 | 0.75 | 0.75 | 5.5 | Moderate |
| *Procyon lotor* | 1 | 1 | 1 | 1 | 0.0 | 1.5 | 5.5 | Moderate |
| *Gulo gulo* | 1 | 1 | 1 | 1 | 0.0 | 0.75 | 4.75 | Moderate |
| *Martes americana* | 0 | 1 | 0 | 1 | 0.75 | 1.5 | 4.25 | Low |
| *Marmota caligata* | 0 | 1 | 1 | 0 | 0.0 | 1.5 | 3.5 | Low |
| *Lepus americanus* | 0 | 0 | 0 | 0 | 1.5 | 1.5 | 3.0 | Low |
| *Neotoma cinerea* | 0 | 0 | 0 | 0 | 0.75 | 0.75 | 1.5 | Low |

**Table S3.** Species within range in each park (JL = Joffre Lakes, GB = Garibaldi), as identified by spatial data provided by the International Union for Conservation of Nature (IUCN) Red List of Threatened Species (IUCN, 2024; https://www.iucnredlist.org/resources/spatial-data-download). We only included terrestrial mammals in range for each park that met our size threshold (average adult body mass >250g as identified by Soria et al., 2021).

| **Species** | **Compiler** | **Year Compiled** | **Citation** | **Park** |
|---|---|---|---|---|
| *Odocoileus hemionus* | IUCN | 2009 | IUCN | JL, GB |
| *Odocoileus virginianus* | IUCN | 2008 | IUCN | JL, GB |
| *Castor canadensis* | IUCN | 2016 | IUCN | JL, GB |
| *Erethizon dorsatum* | IUCN SSC Small Mammal Specialist Group | 2016 | IUCN SSC Small Mammal Specialist Group | JL, GB |
| *Ondatra zibethicus* | IUCN | 2016 | IUCN | JL, GB |
| *Lepus americanus* | IUCN | 2019 | IUCN | JL, GB |
| *Gulo gulo* | IUCN | 2016 | IUCN | JL, GB |
| *Mephitis mephitis* | IUCN | 2016 | IUCN | JL, GB |

| | | | | |
|---|---|---|---|---|
| *Martes americana* | IUCN | 2016 | IUCN | JL, GB |
| *Neovison vison* | IUCN | 2016 | IUCN | JL, GB |
| *Taxidea taxus* | IUCN | 2008 | IUCN | JL |
| *Procyon lotor* | IUCN | 2008 | IUCN | JL, GB |
| *Spilogale gracilis* | IUCN | 2016 | IUCN | JL, GB |
| *Lynx rufus* | IUCN | 2016 | IUCN | JL, GB |
| *Puma concolor* | IUCN | 2015 | IUCN | JL, GB |
| *Lynx canadensis* | Maine Department of Inland Fisheries and Wildlife | 2016 | Maine Department of Inland Fisheries and Wildlife | JL, GB |
| *Ursus americanus* | IUCN | 2016 | IUCN | JL, GB |
| *Neotoma cinerea* | IUCN | 2008 | IUCN | JL, GB |
| *Ursus arctos* | B. McLellan, M Proctor | 2016 | IUCN SSC Bear Specialist Group | JL, GB |
| *Marmota caligata* | IUCN | 2018 | IUCN | JL, GB |
| *Canis latrans* | Hody and Kays 2018 | 2018 | Hody and Kays 2018 | JL, GB |
| *Lontra canadensis* | IUCN | 2015 | IUCN | JL, GB |
| *Vulpes vulpes* | IUCN | 2016 | IUCN | JL, GB |

| | | | | |
|---|---|---|---|---|
| *Oreamnos americanus* | IUCN | 2022 | IUCN | JL, GB |
| *Canis lupus* | IUCN | 2023 | IUCN | JL, GB |

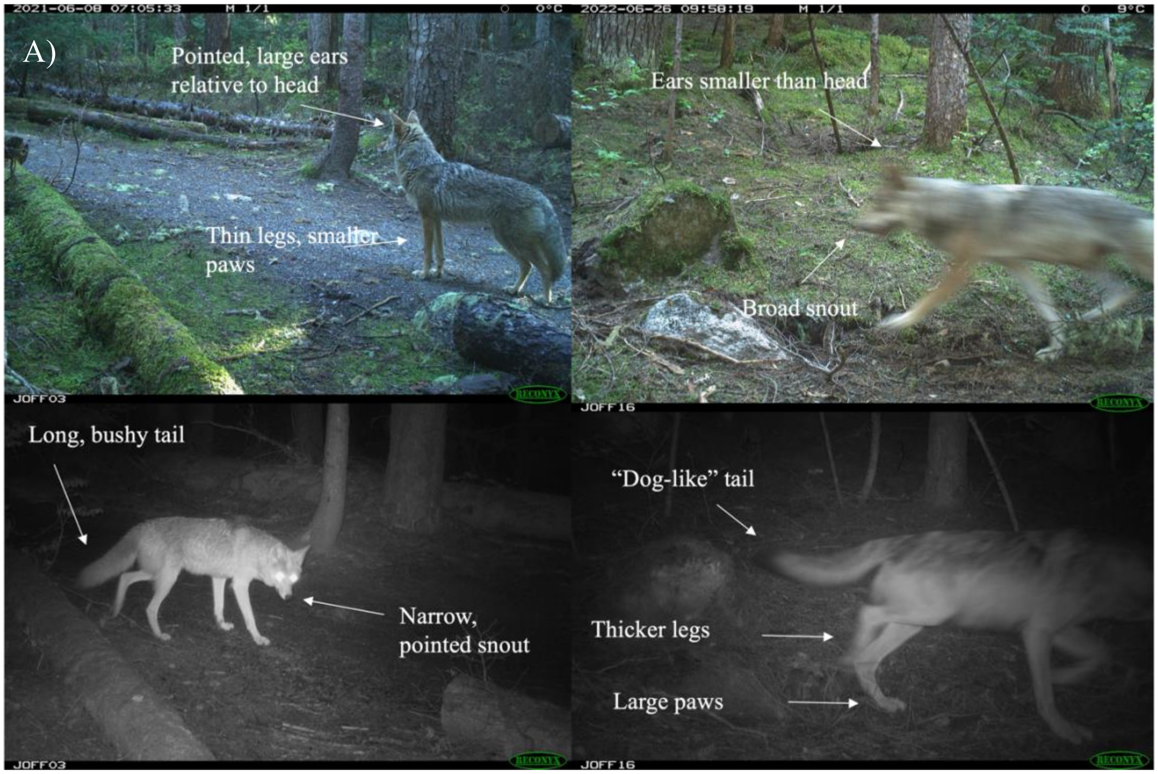

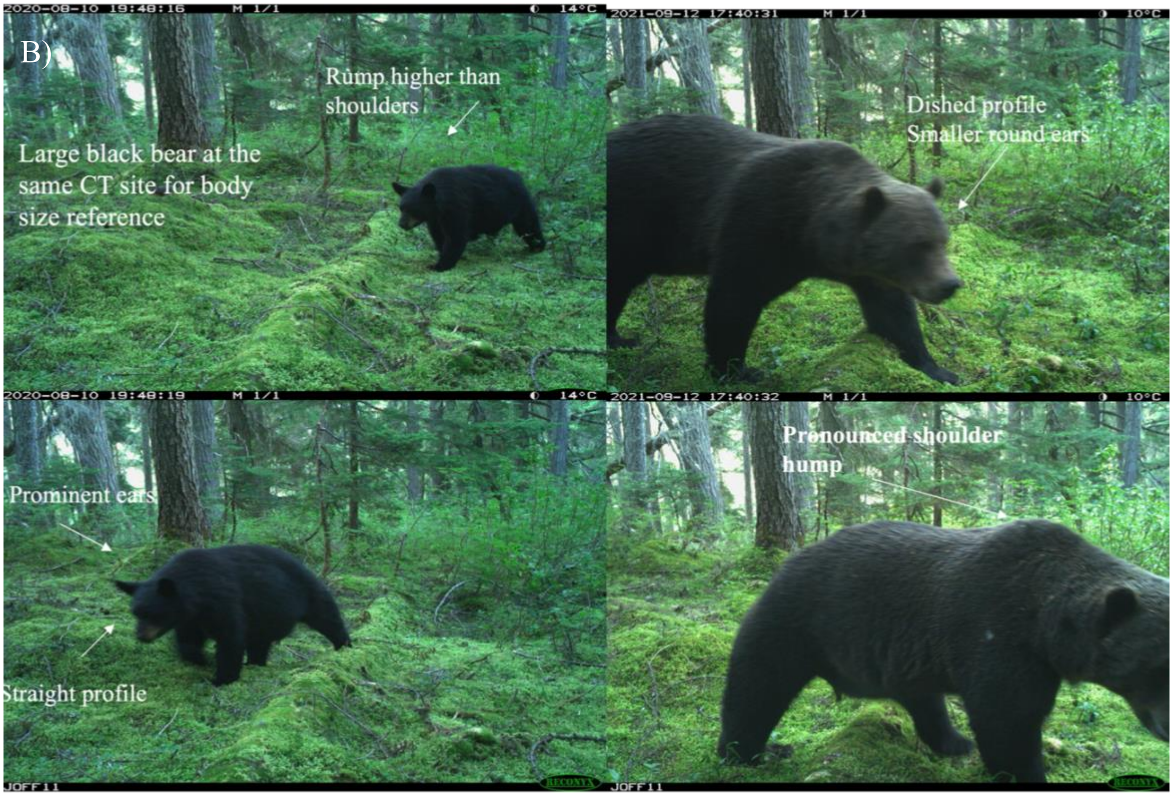
B)
Rump higher than shoulders
Large black bear at the same CT site for body size reference
Dished profile
Smaller round ears
Prominent ears
Straight profile
Pronounced shoulder hump

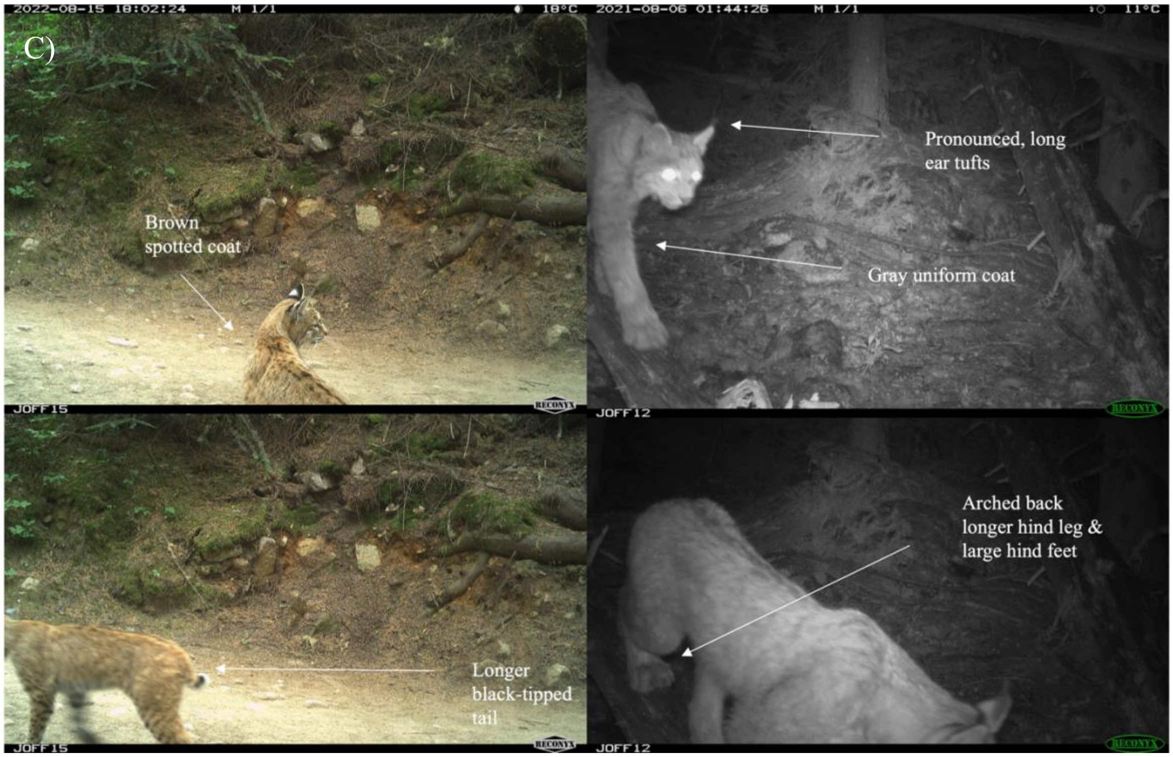
C)
Brown spotted coat
Pronounced, long ear tufts
Gray uniform coat
Arched back longer hind leg & large hind feet
Longer black-tipped tail

**Figure S2.** CT images from our study of rare species (right) detected in Joffre Lakes Park compared to visually similar, more common species (left). Images provide visual confirmation of rarely detected species: A) grey wolf (compared to coyote); B) grizzly bear (compared to black bear); C) Canada lynx (compared to bobcat). Images are annotated with physical traits used to distinguish between species (in white).

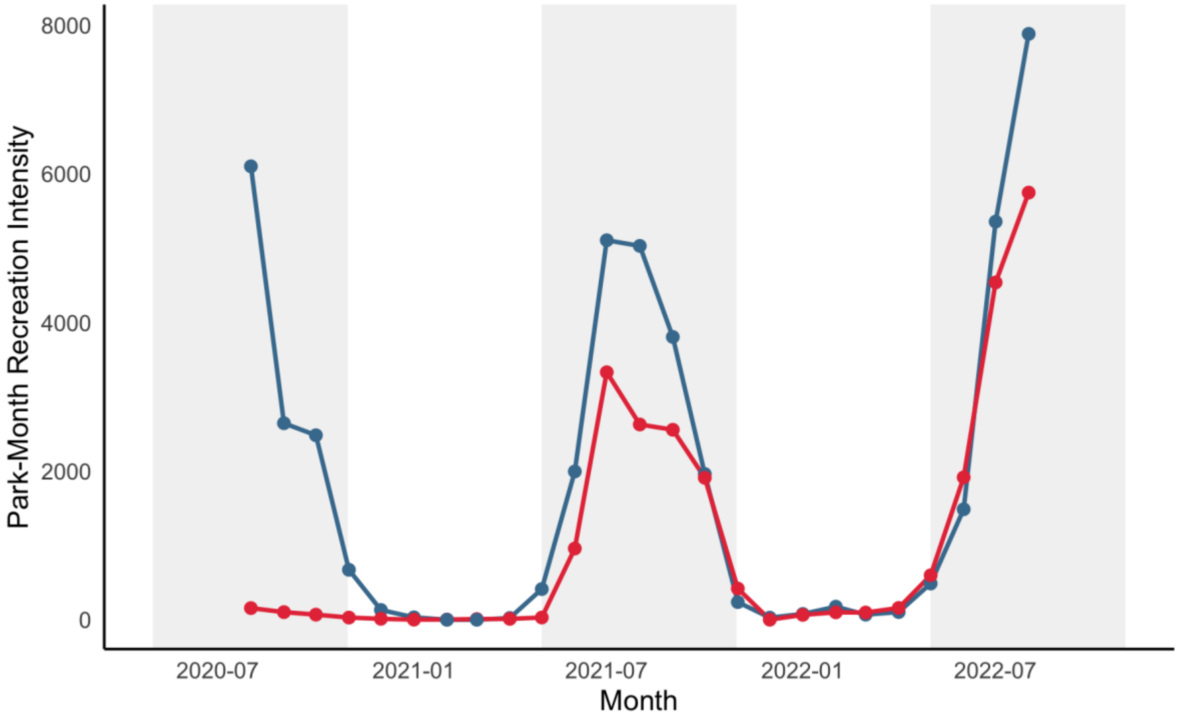


**Figure S3.** Park-Month recreation intensity (independent human detection per 100 CT days) by calendar month in Garibaldi (blue) and Joffre Lakes (red) Parks, Canada. The months included in habitat use models (May-October) are shaded in grey.

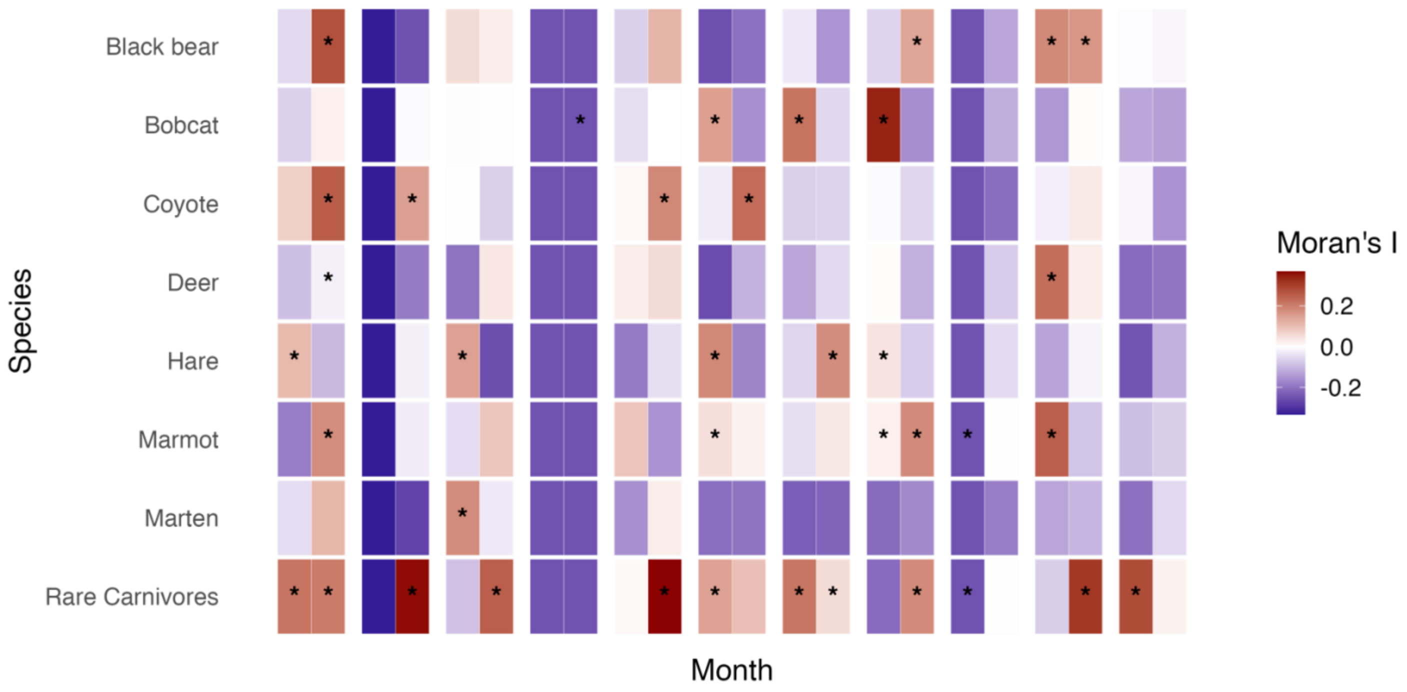


**Figure S4.** Results of a post-hoc Moran's I test for spatial autocorrelation of residuals for species' models. Each square represents a month (28-day period as defined in habitat use model), with Moran's I for Joffre Lakes shown on the right and Garibaldi shown on the left. Moran's I was calculated by species, park, and month using spatially explicit model residuals from Bayesian GLMMs. Spatial relationships were defined using a k-nearest neighbours approach, with each CT site connected to its four nearest neighbours to construct the spatial weights matrix. Positive values (red) indicate spatial clustering of residuals, negative values (purple) indicate spatial dispersion, and values near zero (white) indicate random spatial structure. Asterisks (*) denote months with statistically significant autocorrelation ($p < 0.05$). Spatial autocorrelation of residuals for the rare carnivore group should be interpreted with caution as it is a pooled category. The observed spatial clustering likely reflects the non-overlapping detections of different species, rather than meaningful spatial structure within the group. Thus, despite being statistically significant, this clustering is likely not biologically significant. Overall, we deemed remaining instances of significant autocorrelation to be sufficiently low and thus did not justify modifications to the model structure.

**Table S4.** Results of a post-hoc Durbin-Watson test for temporal autocorrelation of residuals for species' models. Again, the aggregated rare carnivore model results should be interpreted with caution. Values between 1.5 and 2.5 are generally considered acceptable.

| **Species Model** | **Durbin-Watson Statistic** |
|---|---|
| Black bear | 1.78 |
| Bobcat | 2.11 |
| Coyote | 2.13 |
| Deer | 2.33 |
| Hare | 2.06 |
| Marmot | 1.81 |
| Marten | 2.01 |
| Rare Carnivores | 2.35 |

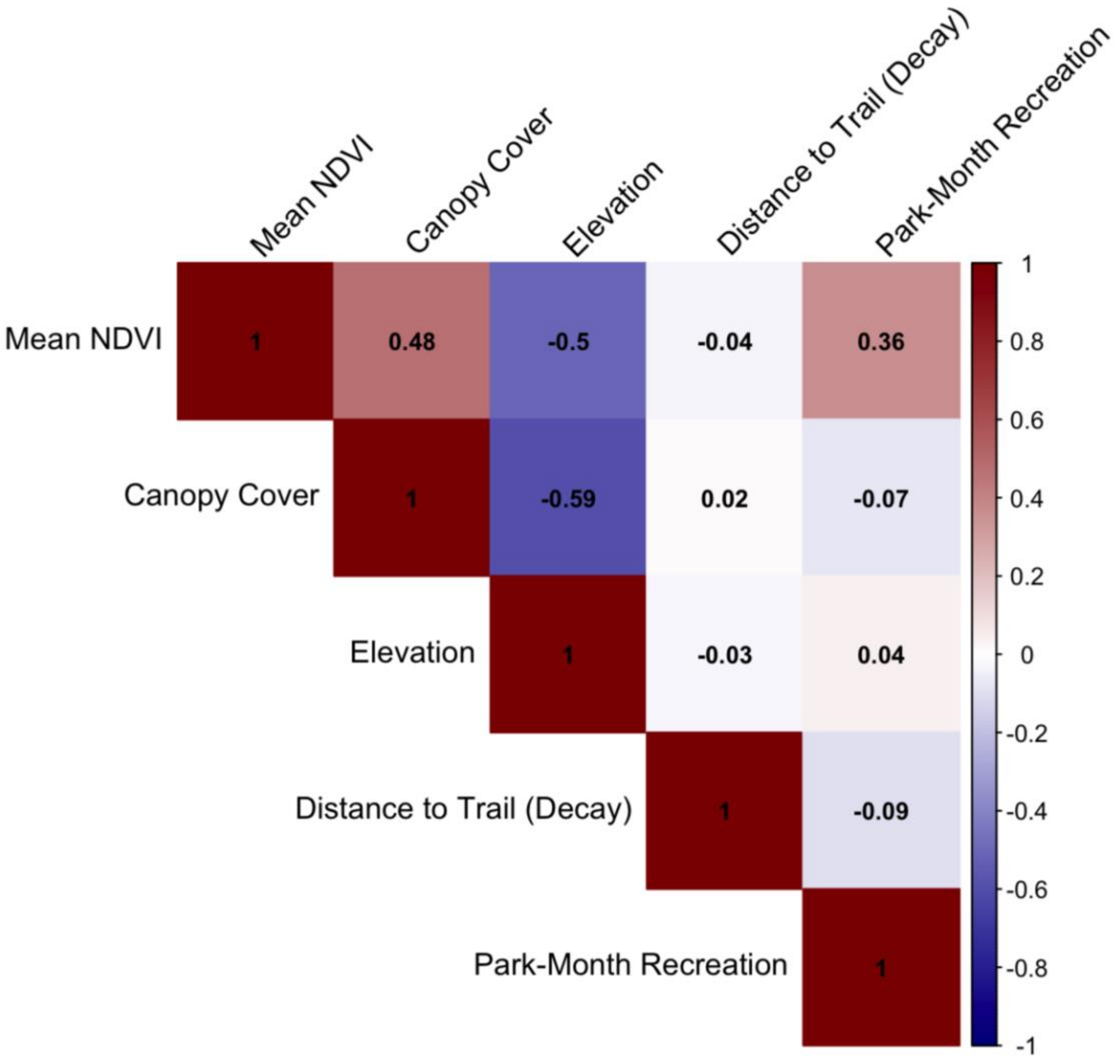


**Figure S5.** Pairwise Pearson correlation coefficient plots for numeric predictor variables used in Bayesian GLMMs. No pair of variables with an absolute correlation coefficient greater than 0.5 (|r| > 0.5) was included in the models (e.g., elevation was omitted).

A)

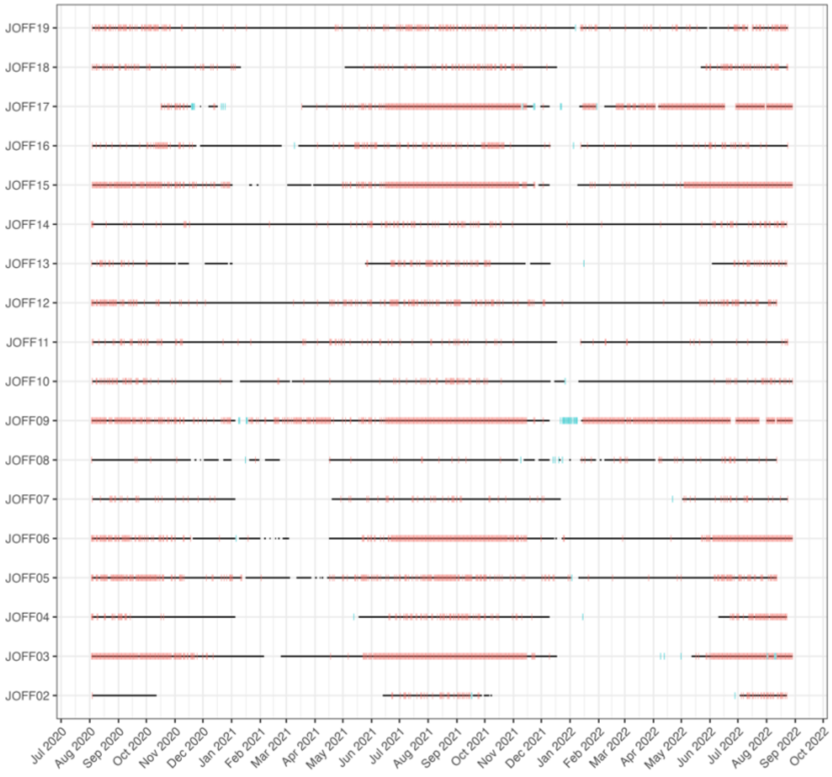
JOFF19
JOFF18
JOFF17
JOFF16
JOFF15
JOFF14
JOFF13
JOFF12
JOFF11
JOFF10
JOFF09
JOFF08
JOFF07
JOFF06
JOFF05
JOFF04
JOFF03
JOFF02
Jul 2020 Aug 2020 Sep 2020 Oct 2020 Nov 2020 Dec 2020 Jan 2021 Feb 2021 Mar 2021 Apr 2021 May 2021 Jun 2021 Jul 2021 Aug 2021 Sep 2021 Oct 2021 Nov 2021 Dec 2021 Jan 2022 Feb 2022 Mar 2022 Apr 2022 May 2022 Jun 2022 Jul 2022 Aug 2022 Sep 2022 Oct 2022

B)

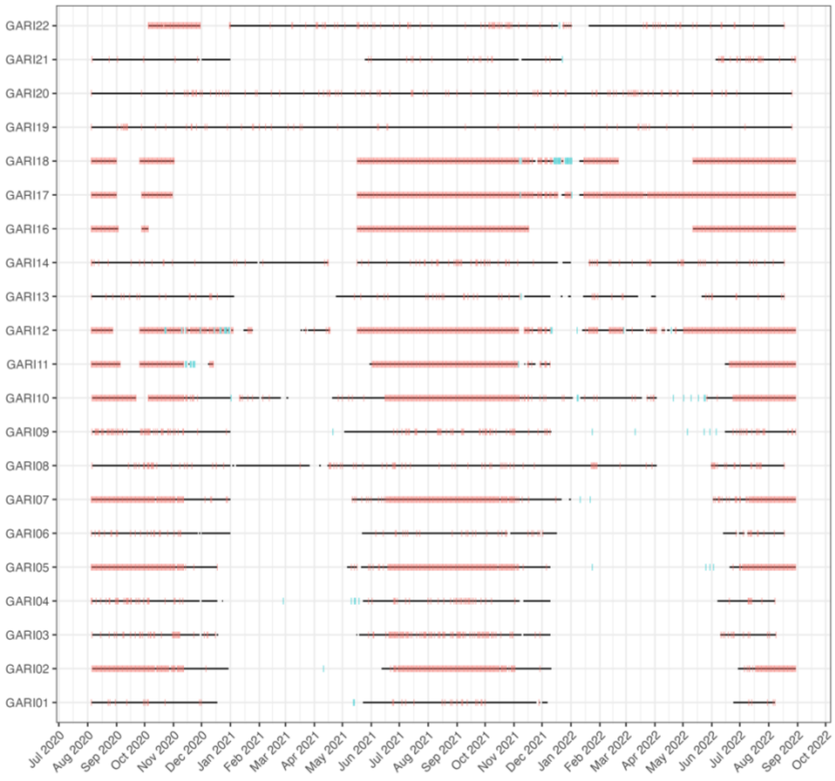
GARI22
GARI21
GARI20
GARI19
GARI18
GARI17
GARI16
GARI14
GARI13
GARI12
GARI11
GARI10
GARI09
GARI08
GARI07
GARI06
GARI05
GARI04
GARI03
GARI02
GARI01
Jul 2020 Aug 2020 Sep 2020 Oct 2020 Nov 2020 Dec 2020 Jan 2021 Feb 2021 Mar 2021 Apr 2021 May 2021 Jun 2021 Jul 2021 Aug 2021 Sep 2021 Oct 2021 Nov 2021 Dec 2021 Jan 2022 Feb 2022 Mar 2022 Apr 2022 May 2022 Jun 2022 Jul 2022 Aug 2022 Sep 2022 Oct 2022

**Figure S6.** Active periods for each CT site in Joffre Lakes (A) and Garibaldi (B). Black lines include dates where a daily timelapse image at 12:00 was taken and unobstructed (e.g., by snow). Pink lines represent dates when a motion-triggered image was taken, and blue lines represent days when we know that the camera view was obstructed. White space represents periods of inactivity for the cameras due to snow cover, camera malfunction, depleted batteries or full storage.

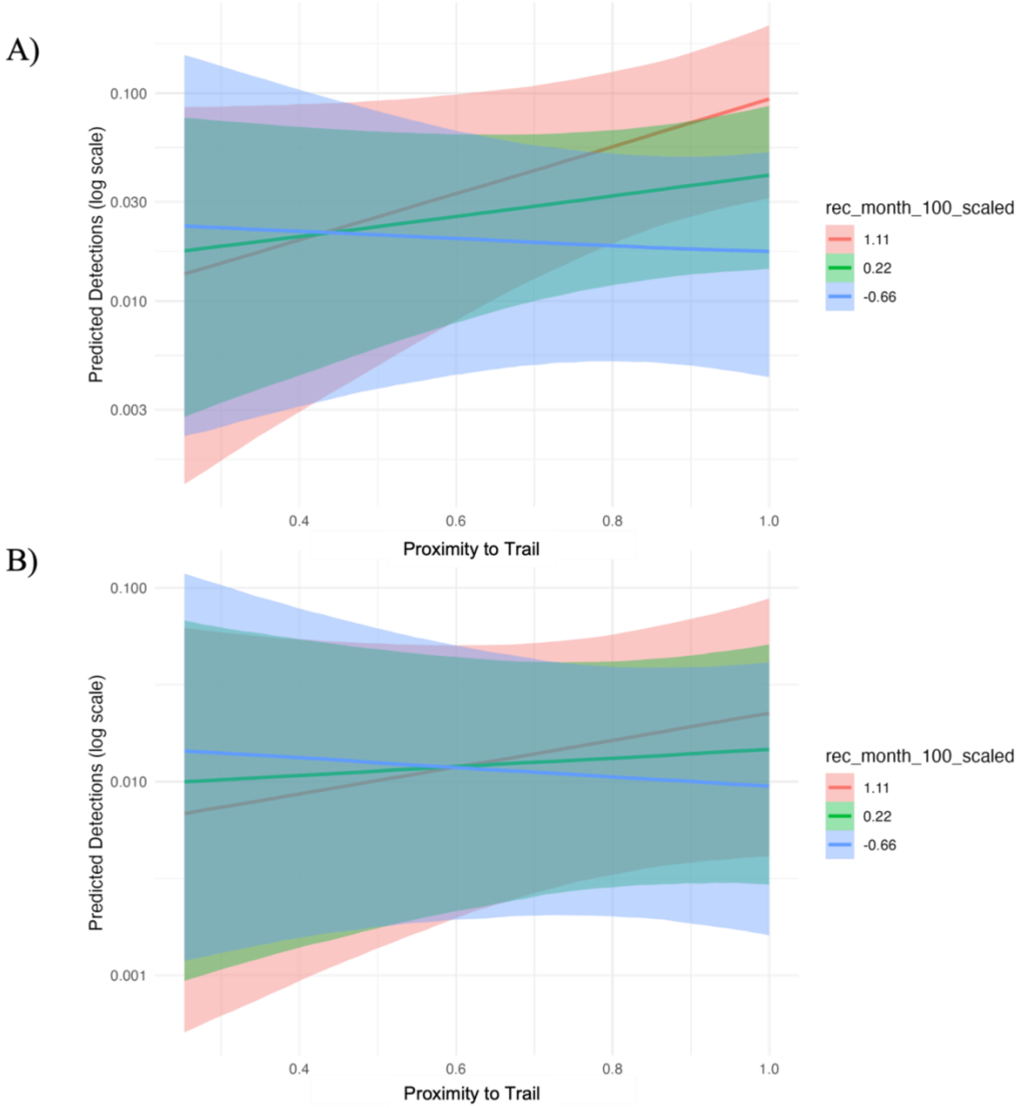


**Figure S7.** Marginal effects plot showing the predicted monthly detection rates (log scale) for bobcats (*Lynx rufus*; A) and marmots (*Marmota caligata*; B) in response to proximity to trail under varying levels of recreational activity. Shaded ribbons represent 95% credible intervals around the posterior median predictions. Line colours vary by recreation intensity level, which correspond to the 1st quartile (low; 0.66), mean (average;0.22), and 3rd quartile (high;1.11) of the recreation intensity variable. All other predictor variables were held at their mean. For both species, the model predicted low baseline detections across all proximity to trail values and wide 95% credible intervals around the predicted values indicate considerable uncertainty in the strength and consistency of this response

**Table S5.** Comparisons of asymptotic estimates of diversity metrics (species richness, Shannon diversity, Pielou's evenness) between Garibaldi (GB) and Joffre Lakes (JL) over time. Shown are point estimates, standard errors (SE), mean differences between groups (Estimate 2 – Estimate 1), 95% confidence intervals (CIs), and associated continuity-corrected p-values derived from bootstrap hypothesis tests. P-values that have had a Benjamini–Hochberg (BH) correction applied are also included to account for multiple comparisons. Significant differences (after BH correction) are bolded (p>0.05).

| **Diversity Metric** | **Comparison** | **Estimate 1** | **SE 1** | **Estimate 2** | **SE 2** | **Mean Difference** | **Lower CI** | **Upper CI** | **P Value** | **BH-Corrected P Value** |
|---|---|---|---|---|---|---|---|---|---|---|
| Species Richness | JL 2020 vs 2021 | 11.98 | 1.71 | 14.96 | 3.45 | 2.75 | -4.85 | 9.89 | 0.45 | 0.61 |
| Species Richness | JL 2020 vs 2022 | 11.98 | 1.71 | 8.99 | 0.72 | -3.02 | -6.39 | 0.62 | 0.1 | 0.31 |
| Species Richness | JL 2021 vs 2022 | 14.96 | 3.45 | 8.99 | 0.72 | -5.77 | -12.68 | 1.3 | 0.11 | 0.31 |
| **Species Richness** | **GB 2020 vs 2021** | **7** | **0.29** | **9** | **0.48** | **2.03** | **0.89** | **3.13** | **0** | **0.02** |
| Species Richness | GB 2020 vs 2022 | 7 | 0.29 | 8.49 | 0.82 | 1.5 | -0.18 | 3.13 | 0.08 | 0.31 |
| Species Richness | GB 2021 vs 2022 | 9 | 0.48 | 8.49 | 0.82 | -0.53 | -2.42 | 1.2 | 0.59 | 0.69 |
| **Species Richness** | **JL vs GB 2020** | **11.98** | **1.71** | **7** | **0.29** | **-5.04** | **-8.36** | **-1.69** | **0.01** | **0.04** |
| Species Richness | JL vs GB 2021 | 14.96 | 3.45 | 9 | 0.48 | -5.75 | -12.44 | 1.42 | 0.11 | 0.31 |
| Species Richness | JL vs GB 2022 | 8.99 | 0.72 | 8.49 | 0.82 | -0.52 | -2.66 | 1.48 | 0.66 | 0.69 |
| Shannon Diversity | JL 2020 vs 2021 | 1.85 | 0.06 | 1.72 | 0.09 | -0.13 | -0.33 | 0.1 | 0.22 | 0.38 |
| **Shannon Diversity** | **JL 2020 vs 2022** | **1.85** | **0.06** | **1.47** | **0.08** | **-0.38** | **-0.59** | **-0.17** | **0** | **0.02** |
| Shannon Diversity | JL 2021 vs 2022 | 1.72 | 0.09 | 1.47 | 0.08 | -0.26 | -0.5 | 0 | 0.05 | 0.27 |

| | | | | | | | | | | |
|---|---|---|---|---|---|---|---|---|---|---|
| Shannon Diversity | GB 2020 vs 2021 | 1.52 | 0.06 | 1.61 | 0.06 | 0.1 | -0.06 | 0.26 | 0.2 | 0.37 |
| Shannon Diversity | GB 2020 vs 2022 | 1.52 | 0.06 | 1.66 | 0.07 | 0.14 | -0.05 | 0.32 | 0.17 | 0.33 |
| Shannon Diversity | GB 2021 vs 2022 | 1.61 | 0.06 | 1.66 | 0.07 | 0.04 | -0.15 | 0.23 | 0.66 | 0.69 |
| **Shannon Diversity** | **JL vs GB 2020** | **1.85** | **0.06** | **1.52** | **0.06** | **-0.33** | **-0.5** | **-0.16** | **0** | **0.02** |
| Shannon Diversity | JL vs GB 2021 | 1.72 | 0.09 | 1.61 | 0.06 | -0.11 | -0.31 | 0.1 | 0.31 | 0.46 |
| Shannon Diversity | JL vs GB 2022 | 1.47 | 0.08 | 1.66 | 0.07 | 0.19 | -0.02 | 0.41 | 0.09 | 0.31 |
| Pielou's Evenness | JL 2020 vs 2021 | 0.74 | 0.05 | 0.63 | 0.06 | -0.11 | -0.27 | 0.04 | 0.15 | 0.31 |
| Pielou's Evenness | JL 2020 vs 2022 | 0.74 | 0.05 | 0.67 | 0.04 | -0.07 | -0.2 | 0.06 | 0.3 | 0.46 |
| Pielou's Evenness | JL 2021 vs 2022 | 0.63 | 0.06 | 0.67 | 0.04 | 0.04 | -0.1 | 0.18 | 0.61 | 0.69 |
| Pielou's Evenness | GB 2020 vs 2021 | 0.78 | 0.03 | 0.73 | 0.03 | -0.05 | -0.14 | 0.04 | 0.35 | 0.49 |
| Pielou's Evenness | GB 2020 vs 2022 | 0.78 | 0.03 | 0.78 | 0.05 | 0 | -0.13 | 0.12 | 0.92 | 0.92 |
| Pielou's Evenness | GB 2021 vs 2022 | 0.73 | 0.03 | 0.78 | 0.05 | 0.04 | -0.07 | 0.16 | 0.5 | 0.64 |
| Pielou's Evenness | JL vs GB 2020 | 0.74 | 0.05 | 0.78 | 0.03 | 0.04 | -0.08 | 0.16 | 0.54 | 0.67 |
| Pielou's Evenness | JL vs GB 2021 | 0.63 | 0.06 | 0.73 | 0.03 | 0.1 | -0.03 | 0.24 | 0.14 | 0.31 |
| Pielou's Evenness | JL vs GB 2022 | 0.67 | 0.04 | 0.78 | 0.05 | 0.1 | -0.03 | 0.24 | 0.14 | 0.31 |

**Table S6.** Posterior estimates and 95% credible intervals for the standard deviation of the random intercept (CT site) for each species model. Values represent the estimated variability in baseline detection rates across CT sites after accounting for fixed effects.

| Species Model | Estimate | Lower 95% CI | Upper 95% CI | Rhat |
|---|---|---|---|---|
| Black bear | 1.94 | 1.23 | 2.92 | 1.00 |
| Bobcat | 0.69 | 0.04 | 1.73 | 1.00 |
| Coyote | 0.9 | 0.06 | 2.1 | 1.00 |
| Deer | 1.22 | 0.76 | 1.79 | 1.00 |
| Hare | 1.91 | 1.31 | 2.76 | 1.00 |

| | | | | |
|---|---|---|---|---|
| Marmot | 2.16 | 1.25 | 3.53 | 1.00 |
| Marten | 0.72 | 0.09 | 1.42 | 1.00 |
| Rare Carnivores | 0.68 | 0.03 | 1.89 | 1.00 |

**Table S7.** Full results of the Bayesian GLMMs estimating the effect of recreation on habitat use while controlling for other sources of variation, by species. Lower and upper CI are the lower and upper 80 and 95% credible intervals (CIs) for each posterior estimate. Bolded rows represent posterior estimates with 80 and or 95% CIs that do not span 0. mean_ndvi_scaled = Mean Normalized Difference in Vegetation Index per month at each CT site, measured in 16-day intervals (250-meter resolution). Mean is the average of each 16-day value which overlaps a minimum of 5 days with a given month. canopy_cover_scaled = percentage of vegetation taller than 2m (100m radius around CT site) from National Terrestrial Ecosystem Monitoring System (NTEMS*)*. rec_month_100_scaled = Monthly total independent human detections across all CT sites within each park per 100 CT days. prox_to_trail_decay = distance in meters of a CT site to the nearest hiking trail with an exponential decay function (-distance/ 500m).

| Species Model | Variable | Estimate | Lower 95% CI | Upper 95% CI | Lower 80% CI | Upper 80% CI | Rhat |
|---|---|---|---|---|---|---|---|
| Black bear | Intercept | -4.33 | -6.41 | -2.45 | -5.64 | -3.09 | 1 |
| | mean_ndvi_scaled | **0.38** | -0.16 | 0.96 | **0.02** | **0.75** | 1 |
| | canopy_cover_scaled | -0.12 | -0.92 | 0.72 | -0.63 | 0.41 | 1 |
| | rec_month_100_scaled | **-0.81** | -2.03 | 0.36 | **-1.6** | **-0.03** | 1 |
| | prox_to_trail_decay | -0.9 | -3.17 | 1.54 | -2.4 | 0.63 | 1 |
| | rec_month_100_scaled*prox_to_trail_decay | 0.14 | -1.28 | 1.57 | -0.8 | 1.07 | 1 |
| Bobcat | Intercept | -7.6 | -10.01 | -5.5 | -9.1 | -6.16 | 1 |
| | mean_ndvi_scaled | 0.48 | -0.46 | 1.55 | -0.16 | 1.15 | 1 |
| | canopy_cover_scaled | 0.07 | -0.72 | 0.87 | -0.44 | 0.58 | 1 |
| | rec_month_100_scaled | -0.73 | -2.65 | 1.23 | -1.99 | 0.54 | 1 |
| | prox_to_trail_decay | 0.77 | -1.59 | 3.26 | -0.8 | 2.36 | 1 |
| | rec_month_100_scaled*prox_to_trail_decay | **1.69** | -0.4 | 3.76 | **0.33** | **3.03** | 1 |
| Coyote | Intercept | -6.06 | -8.53 | -3.83 | -7.59 | -4.58 | 1 |
| | mean_ndvi_scaled | 0.16 | -0.75 | 1.11 | -0.42 | 0.76 | 1 |
| | canopy_cover_scaled | -0.42 | -1.37 | 0.42 | -1 | 0.13 | 1 |

| | | | | | | | |
|---|---|---|---|---|---|---|---|
| | rec_month_100_scaled | 0.39 | -1.55 | 2.31 | -0.88 | 1.64 | 1 |
| | prox_to_trail_decay | -0.3 | -2.74 | 2.3 | -1.9 | 1.34 | 1 |
| | rec_month_100_scaled*prox_to_trail_decay | -1.38 | -3.64 | 0.82 | -2.86 | 0.07 | 1 |
| Deer | Intercept | -3.26 | -4.71 | -1.97 | -4.16 | -2.39 | 1 |
| | mean_ndvi_scaled | **0.67** | **0.27** | **1.09** | **0.41** | **0.94** | 1 |
| | canopy_cover_scaled | -0.2 | -0.7 | 0.32 | -0.52 | 0.14 | 1 |
| | rec_month_100_scaled | **-0.64** | -1.57 | 0.29 | **-1.24** | **-0.04** | 1 |
| | prox_to_trail_decay | -1.03 | -2.64 | 0.7 | -2.08 | 0.05 | 1 |
| | rec_month_100_scaled*prox_to_trail_decay | 0.49 | -0.6 | 1.56 | -0.21 | 1.18 | 1 |
| Hare | Intercept | -4.58 | -6.54 | -2.75 | -5.82 | -3.37 | 1 |
| | mean_ndvi_scaled | 0.07 | -0.4 | 0.54 | -0.24 | 0.38 | 1 |
| | canopy_cover_scaled | **0.96** | **0.14** | **1.88** | **0.42** | **1.53** | 1 |
| | rec_month_100_scaled | 0.66 | -0.45 | 1.78 | -0.06 | 1.39 | 1 |
| | prox_to_trail_decay | 0 | -2.24 | 2.28 | -1.45 | 1.47 | 1 |
| | rec_month_100_scaled*prox_to_trail_decay | -0.66 | -1.91 | 0.58 | -1.48 | 0.15 | 1 |
| Marmot | Intercept | -7.75 | -10.67 | -5.2 | -9.58 | -6.02 | 1 |
| | mean_ndvi_scaled | -0.2 | -1.04 | 0.61 | -0.73 | 0.32 | 1 |
| | canopy_cover_scaled | **-1.03** | **-2.15** | **-0.01** | **-1.72** | **-0.35** | 1 |
| | rec_month_100_scaled | -0.73 | -2.22 | 0.7 | -1.67 | 0.21 | 1 |
| | prox_to_trail_decay | 0.27 | -2.55 | 3.26 | -1.59 | 2.16 | 1 |
| | rec_month_100_scaled*prox_to_trail_decay | **1.22** | -0.5 | 2.96 | **0.1** | **2.33** | 1 |
| Marten | Intercept | -5.97 | -7.57 | -4.55 | -6.95 | -5.02 | 1 |
| | mean_ndvi_scaled | 0.13 | -0.3 | 0.55 | -0.15 | 0.41 | 1 |
| | canopy_cover_scaled | **-0.55** | **-1.04** | **-0.09** | **-0.85** | **-0.26** | 1 |
| | rec_month_100_scaled | **-1.1** | -2.5 | 0.26 | **-2** | **-0.21** | 1 |
| | prox_to_trail_decay | **1.09** | -0.5 | 2.81 | **0.05** | **2.16** | 1 |
| | rec_month_100_scaled*prox_to_trail_decay | 0.33 | -1.19 | 1.87 | -0.66 | 1.34 | 1 |
| Rare carnivores | Intercept | -6.72 | -9.01 | -4.72 | -8.15 | -5.36 | 1 |
| | mean_ndvi_scaled | **-0.57** | -1.39 | 0.25 | **-1.1** | **-0.04** | 1 |
| | canopy_cover_scaled | -0.44 | -1.22 | 0.34 | -0.94 | 0.06 | 1 |
| | rec_month_100_scaled | 0.23 | -1.71 | 2.16 | -1.02 | 1.48 | 1 |
| | prox_to_trail_decay | -0.79 | -3.17 | 1.68 | -2.36 | 0.8 | 1 |
| | rec_month_100_scaled*prox_to_trail_decay | -0.57 | -2.83 | 1.7 | -2.04 | 0.9 | 1 |

**Table S8.** Bayesian p-values for each species model, calculated from posterior predictive checks based on the mean of the observed response variable. For each model, 1,000 replicated datasets were drawn from the posterior predictive distribution, and the proportion of simulated means greater than or equal to the observed mean was used to calculate the *p*-value. Values near 0.5 indicate good model fit (Gelman et al., 1996).

| **Model** | **Bayesian p-value** |
|---|---|
| Black bear | 0.64 |
| Bobcat | 0.59 |
| Coyote | 0.71 |
| Deer | 0.64 |
| Hare | 0.61 |
| Marmot | 0.59 |
| Marten | 0.55 |
| Rare Carnivores | 0.66 |

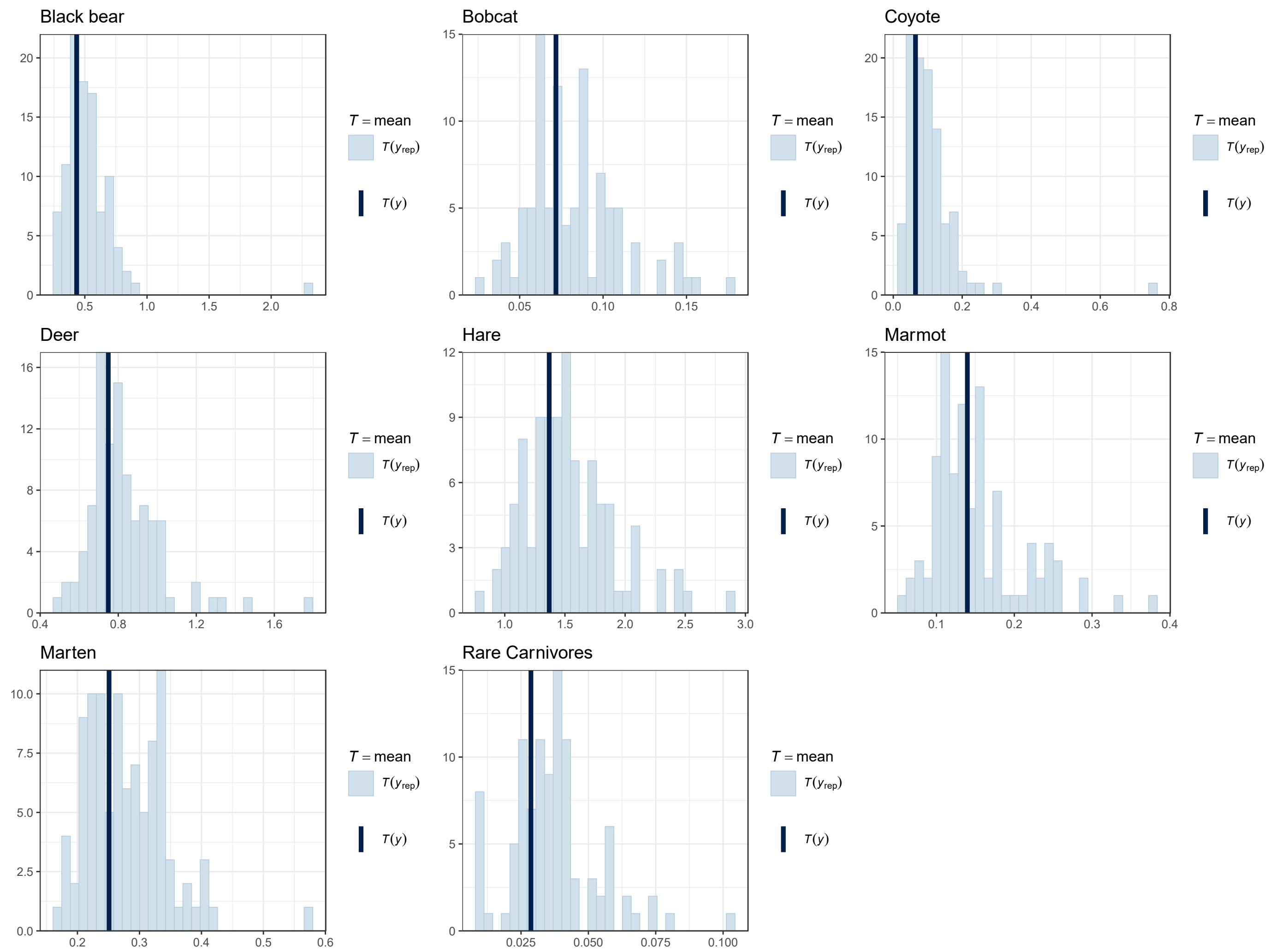


**Figure S8.** Posterior predictive check plots for each species model. Each plot compares the observed mean detection rate to the distribution of simulated means from 1,000 posterior predictive draws. T(*yrep*) (light blue) is the distribution of the simulated means and T(*y*) (dark blue) is the observed mean of the actual data.

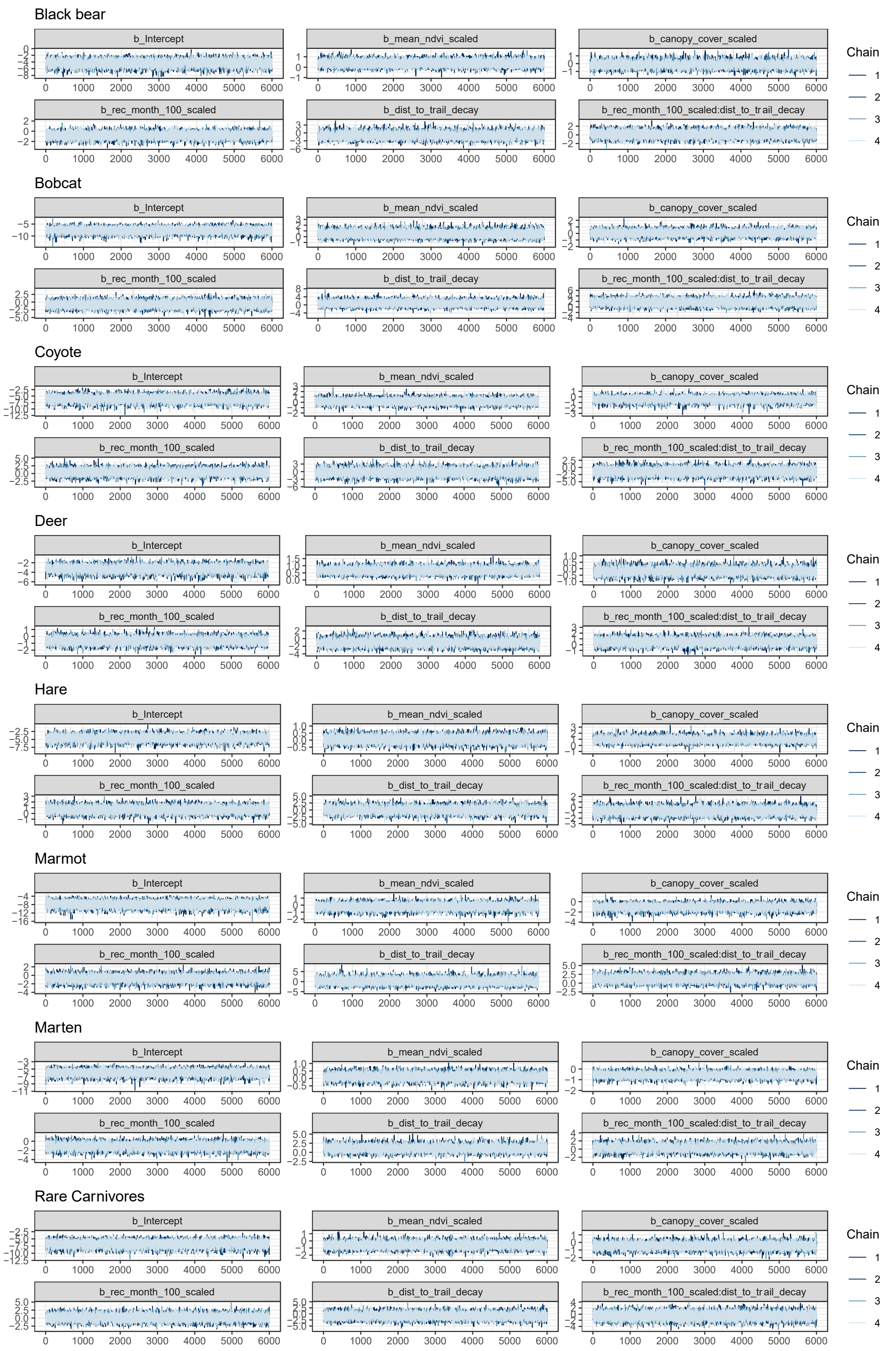
Black bear
Bobcat
Coyote
Deer
Hare
Marmot
Marten
Rare Carnivores
b_Intercept
b_mean_ndvi_scaled
b_canopy_cover_scaled
b_rec_month_100_scaled
b_dist_to_trail_decay
b_rec_month_100_scaled:dist_to_trail_decay
Chain
1
2
3
4

**Figure S9.** Trace plots for the fixed effects in each species model. Each plot shows the Markov Chain Monte Carlo (MCMC) samples across four chains to assess sampling behaviour and convergence for the posterior distributions. *dist_to_trail_decay* corresponds to *prox_trail_decay*.

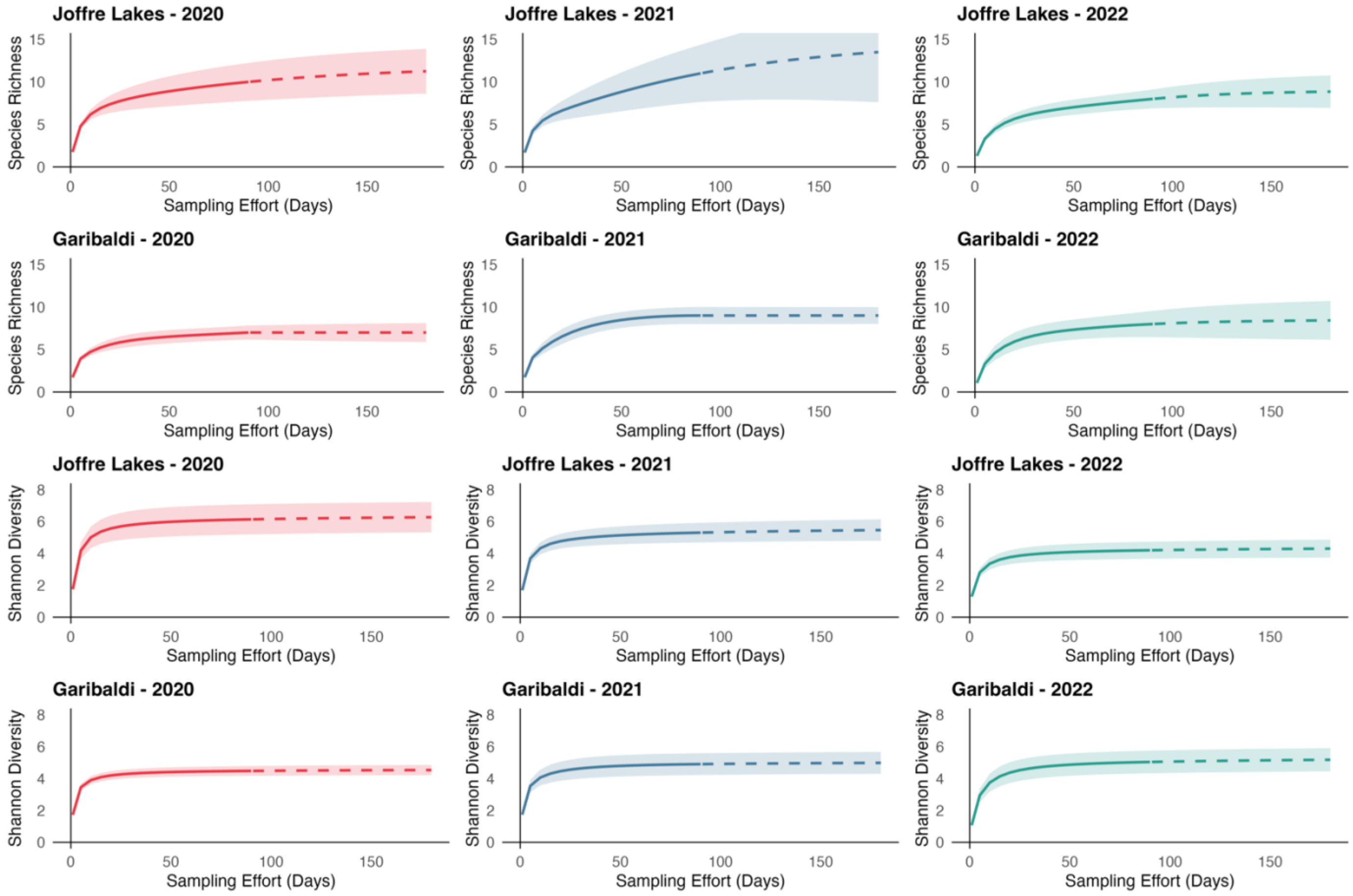


**Figure S10.** Species accumulation curves for species richness (q = 0), and Shannon diversity (q = 1), by year and park. Year is a 90-day period in the spring-fall of the corresponding year: 2020 (red), 2021 (blue), and 2022 (green). Solid lines represent the interpolation of the associated metric, based on CT detections. The dashed lines represent the extrapolated model-based predictions. Shading around the lines represents the 95% confidence intervals.
